# *Clostridioides difficile* toxin A and toxin B inhibit toxin-specific adaptive immune responses through glucosyltransferase-dependent activity

**DOI:** 10.1101/2025.07.30.667646

**Authors:** Jeffrey R. Maslanka, Jennifer A. Londregan, Joshua E. Denny, Ellie N. Hulit, Nontokozo V. Mdluli, F. Christopher Peritore-Galve, Md Zahidul Alam, Mohamad-Gabriel Alameh, D. Borden Lacy, Joseph P. Zackular, Michael C. Abt

## Abstract

*Clostridioides difficile* colonizes the gastrointestinal tract and secretes two virulence factors: toxin A (TcdA) and toxin B (TcdB). Protective immunity against *C. difficile* infection is limited as patients are susceptible to multiple rounds of recurrent infections. The factors determining whether immunity to TcdA and TcdB is generated remain incompletely defined. We determined that *C. difficile*-infected mice generate antibody and IL-17A-producing CD4^+^ T cell responses to TcdA, but not TcdB. To determine the mechanism of the failed anti-TcdB immunity, *C. difficile* mutant strains expressing glucosyltransferase inactive (GTX) TcdA, and/or glucosyltransferase inactive TcdB were used. Infection with TcdB_GTX_ or dual mutant (TcdA_GTX_ TcdB_GTX_) restored TcdB-specific antibody responses, while infection with TcdA_GTX_ or TcdA_GTX_ TcdB_GTX_ led to an earlier induction of TcdA-specific antibodies. Finally, infection with the dual GTX mutant enhanced TcdA and TcdB-specific CD4^+^ T cell responses. These data demonstrate that the glucosyltransferase activity of TcdA and TcdB hinders the antigen-specific adaptive immune response to itself and may be a mechanism that underlies high recurrence rates following *C. difficile* infection in patients.

## Introduction

The nosocomial pathogen *Clostridioides difficile* represents a major healthcare challenge partly due to the prevalence of recurrent infections. One in four *C. difficile* patients experience recurrent disease, with up to 50% of recurrent infections being caused by the same strain as the primary infection^1–3^. The risk of disease recurrence increases with each episode^3^, leading to a debilitating quality of life and disruption of medical treatment. The greatest risk for *C. difficile* recurrence occurs within 24 days of the initial onset of symptoms^4^. The high rate of recurrence following *C. difficile* infection coupled with the timing of recurrent disease strongly suggest that the quality and protective capacity of the adaptive immune response generated following primary *C. difficile* infection may be significantly compromised.

*C. difficile* pathogenesis is mediated through two exotoxins, toxin A (TcdA) and toxin B (TcdB). These homologous proteins share 63% sequence similarity and have conserved effector function^5^. TcdA and TcdB are GTPase inhibitors that glucosylate host Rho-family GTPases, resulting in a loss of function within the host cell^6^. At steady state, GTPase enzymes hydrolyze GTP to perform many critical cell functions such as regulation of the actin cytoskeleton and cell cycle progression^7^. Cellular intoxication by TcdA and TcdB leads to breakdown of the actin cytoskeleton, and inhibition of key cellular processes eventually leading to cell death^8^. Toxin virulence is primarily caused by the glucosyltransferase activity of TcdA and TcdB, as point mutations in the glucosyltransferase domain of TcdA and TcdB renders *C. difficile* asymptomatic *in vivo*^9, 10^. Although both toxins are capable of causing cell and tissue damage, TcdB is required for symptomatic disease while TcdA is dispensable in human disease and animal models^11–14^.

In humans, high serum antibody responses to both *C. difficile* toxins correlate with lower rates of recurrent disease overall^15–22^. Moreover, data from clinical trials demonstrate that a TcdB-specific monoclonal antibody (Bezlotuxumab), but not a TcdA-specific monoclonal antibody (Actoxumab), significantly reduced recurrence rates^23^. These data suggest that TcdB- specific antibodies can prevent symptomatic disease recurrence, while TcdA-specific antibodies may be dispensable^23^. Based on these reports, we hypothesize that a subset of patients fail to generate anti-toxin antibody responses and are thus more likely to develop a recurrent infection. Mouse models of *C. difficile* reveal a lack of protection from secondary infection, and this lack of protection is associated with an absence of serum antibodies to TcdB^24, 25^. Vaccine studies in mice targeting TcdA and TcdB demonstrate that robust protective immunity can be generated following vaccination^24–31^, however, these protective responses are not elicited *in vivo* following infection.

The generation of adaptive immune response to *C. difficile* toxins following natural infection has not been extensively explored and it is not clear why *C. difficile*-specific immunity fails to develop following infection. Further, there are no studies investigating the mucosal toxin-specific CD4^+^ T cell response generated following infection. In this study, we demonstrate that mice generate robust and neutralizing antibody responses to TcdA following infection but fail to generate significant antibody responses to TcdB. Additionally, TcdA-responsive CD4^+^ T cells are generated following infection; however, these mice do not generate TcdB-responsive CD4^+^ T cells. Infection with toxin mutant strains that express intact, but enzymatically inactive toxin (glucosyltransferase mutant) leads to earlier and more robust antibody and CD4^+^ T cell responses to TcdA and rescues antibody and CD4^+^ T cell responses to TcdB. These data suggest that the glucosyltransferase activity of TcdA and TcdB inhibit formation of toxin-specific adaptive immune response and identifies a novel mechanism of immune evasion by *C. difficile* toxins.

## Results

### Mice generate robust antibody responses to TcdA but fail to generate TcdB-specific antibody responses following infection

To investigate adaptive immune responses to *C. difficile* toxins following infection, we utilized a well-defined mouse model of *C. difficile* infection (**Supplemental Figure 1A**)^32–35^. Antibiotic-treated mice were infected with *C. difficile* spores (CD196 strain), experienced acute disease (**Figure 1A,B**), and were chronically infected with toxin-producing *C. difficile* for weeks following infection (**Supplemental Figure 1B,C**). *C. difficile*-infected (CDI) or antibiotic-treated (ABX) control mice were sacrificed on days 7, 14, 21, and 35; timepoints to define the kinetics and magnitude of the humoral and cellular adaptive immune response to *C. difficile* toxins following infection.

**Figure 1.**
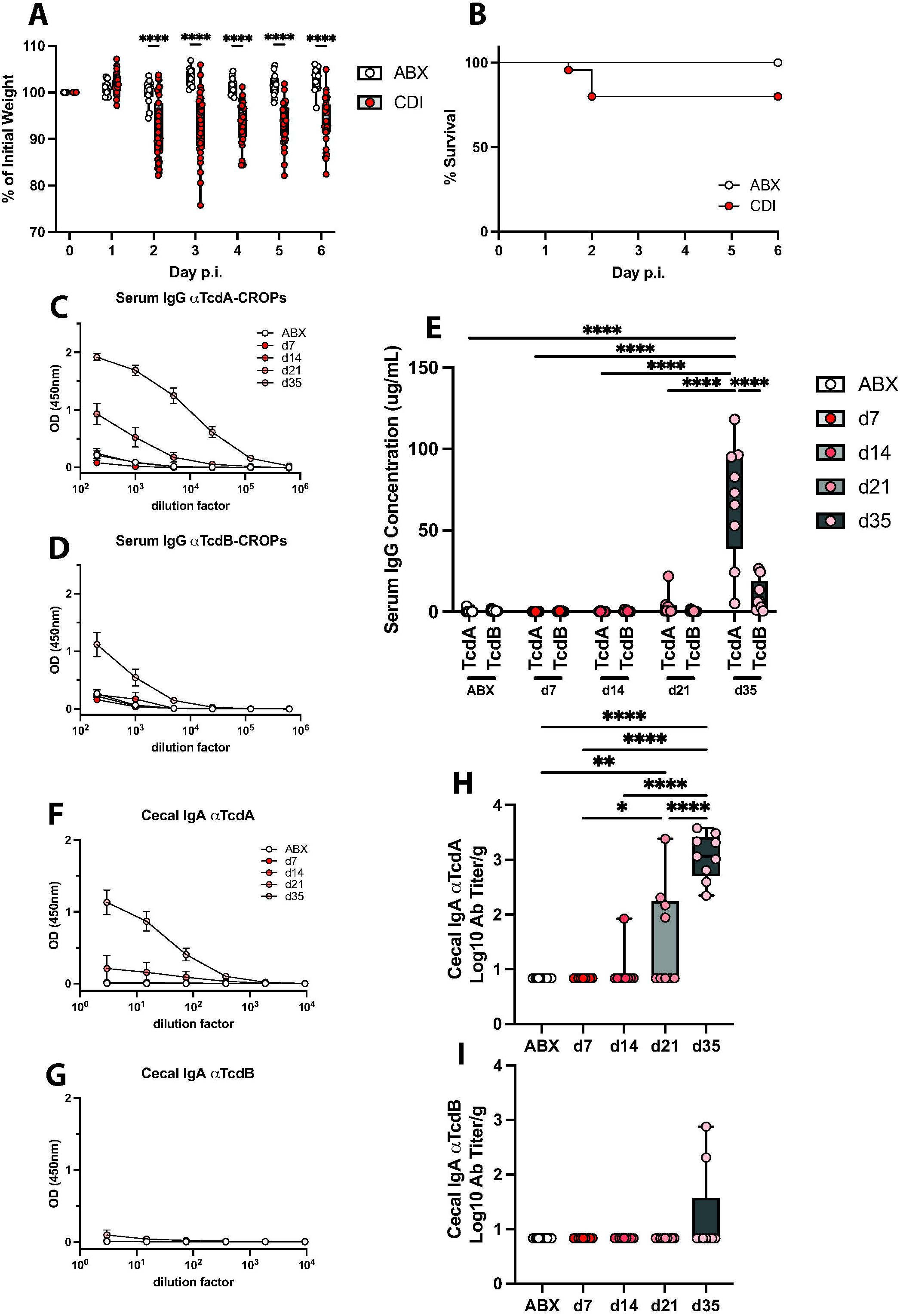
Mice generate robust toxin-specific antibody responses to TcdA but not TcdB following *C. difficile* infection. (A) Weight loss and (B) mortality following *C. difficile* infection. (C) Optical density of serum IgG specific for TcdA-CROPs or (D) TcdB-CROPs at various timepoints post *C. difficile* infection. (E) Quantification of serum IgG specific for TcdA-CROPs and TcdB-CROPs. (F) Optical density of cecal IgA specific for TcdA-CROPs or (G) TcdB-CROPs. (H) IgA titer per gram of cecal content specific for TcdA-CROPs or (I) TcdB-CROPs. Representative data were pooled from two independent experiments. ABX n=14 and *C. difficile*-infected n=9 per timepoint. (A,H,I) One-way ANOVA with Tukey’s multiple comparison test. (E) Two-way ANOVA with Tukey’s multiple comparison test. Statistical significance is indicated as follows: *** P < 0.001; **** P < 0.0001.

Serum IgG responses to the combined repetitive oligopeptides (CROPs) domain of TcdA and TcdB were measured following infection as this region is immunodominant^26, 36, 37^. *C. difficile*-infected mice had TcdA-specific IgG that was detected on day 21 and peaked on day 35 post-infection (**Figure 1C**), however, only low levels of TcdB-specific IgG were detected after 35 days of infection (**Figure 1B**). The IgG response to TcdA and TcdB were quantified and directly compared by using mouse monoclonal antibodies against TcdA and TcdB as a standard. There was a robust TcdA-specific IgG response in the serum at day 35 post-infection, with no significant induction of TcdB-specific IgG compared to antibiotic-treated control mice (TcdB: ABX vs d35, p=0.3765) (**Figure 1E**). To confirm that the antibody response to the CROP domain was comprehensive of the total anti-toxin antibody response, serum IgG to full-length TcdA and TcdB was determined. There was a significant IgG response to full-length TcdA in the serum at day 35 post-infection, but no significant IgG response to full-length TcdB, ruling out the presence of IgG specific to other domains of TcdB (**Supplemental Figure 2A-C**). Therefore, TcdA-specific IgG was induced at day 35 post-infection and was significantly greater than the TcdB-specific IgG response at this time point. Next, since *C. difficile* is an intestinal luminal pathogen, TcdA- and TcdB-specific IgA in the cecal content was assessed. A portion of mice infected with *C. difficile* had detectable TcdA-specific IgA at day 21 post infection while all mice had high titers of TcdA-specific IgA by day 35 post infection (**Figure 1F,H**). However, there was no significant TcdB-specific IgA response in the cecal content compared to antibiotic-treated control mice (**Figure 1G,I**), mirroring the kinetics of the serum IgG response.

To determine if anti-toxin antibody responses elicited following infection were protective, an *in vitro* toxin neutralization assay was established. Vero cells were used to assess active *C. difficile* toxin by cell rounding (**Figure 2A**). TcdA and TcdB were tittered to the lowest concentrations that reproducibly rounded all Vero cells (data not shown). Actoxumab and Bezlotoxumab were used as positive controls to demonstrate neutralization capacity of TcdA and TcdB *in vitro* (**Supplemental Figure 3**). Toxin neutralization was measured using serum from *C. difficile*-infected or antibiotic-treated control mice at indicated timepoints. Serum from antibiotic-treated control mice did not prevent cell rounding from TcdA or TcdB challenge, whereas serum from *C. difficile*-infected mice at day 35 post-infection was able to protect cells from TcdA induced cell rounding (**Figure 2A,B**) but not TcdB induced cell rounding (**Figure 2A,C**). These data functionally validate our anti-toxin antibody data, confirming that the antibody response generated to TcdA is neutralizing, while mice fail to generate neutralizing antibody responses to TcdB.

**Figure 2.**
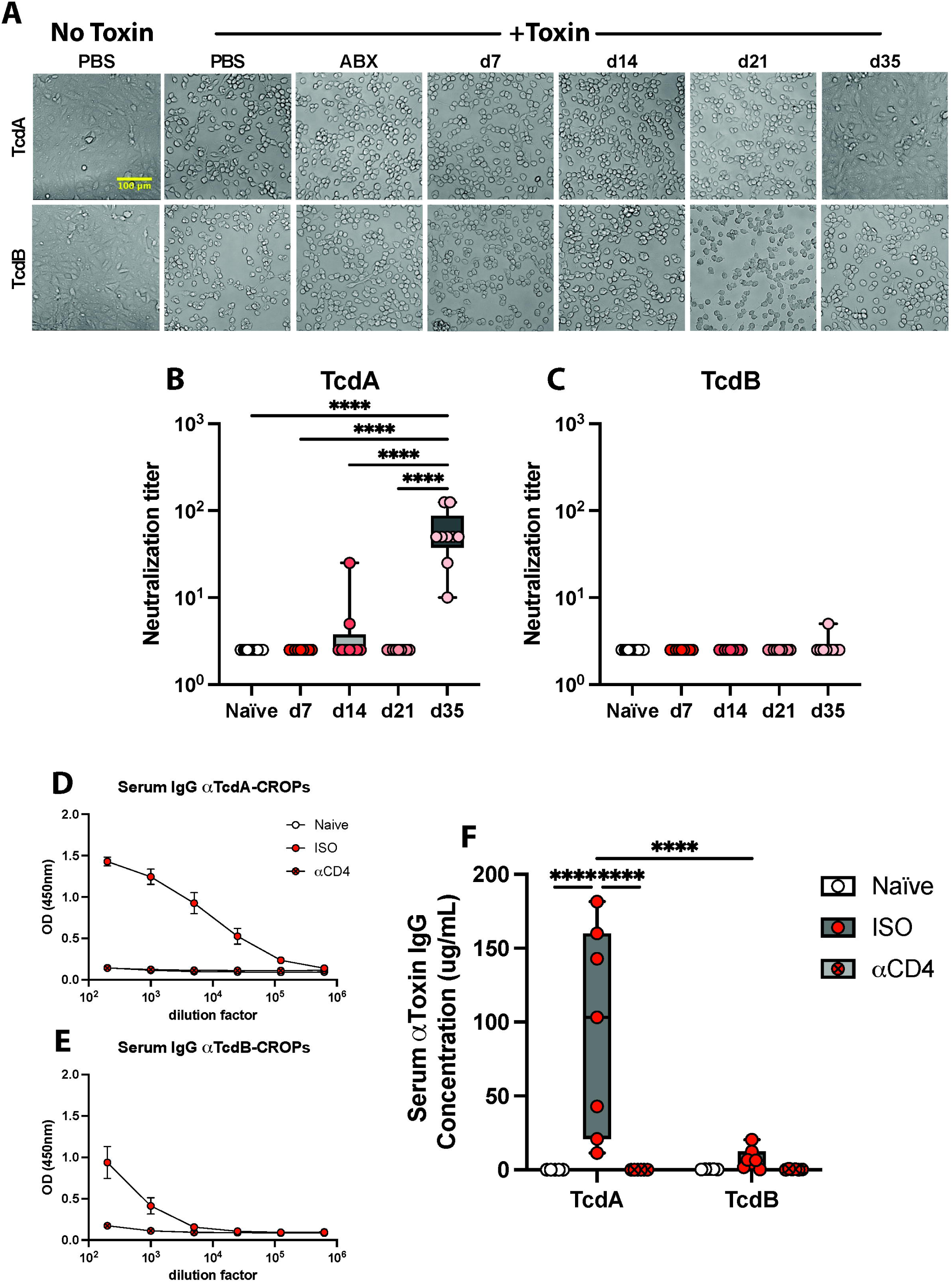
Serum antibody responses to TcdA are neutralizing and CD4^+^ T cell dependent. (A) Representative brightfield images of Vero cells incubated the presence of TcdA and TcdB with and without serum isolated at various timepoints post-infection. (B,C) Quantification of serum neutralization titer determined from various timepoints post-infection. Representative data were pooled from two independent experiments. (D) The optical density of serum IgG specific for TcdA or (E) TcdB at day 35 post *C. difficile* infection. Infected mice were treated with multiple doses of an isotype control antibody or a CD4 depletion antibody. (A-C) ABX n=14 and *C. difficile*-infected n=9 per timepoint. One-way ANOVA with Tukey’s multiple comparison test. Statistical significance is indicated as follows: **** P < 0.0001. (D-F) Quantification of serum IgG specific for TcdA-CROP and TcdB-CROP. Naïve n=5, *C. difficile*-infected isotype treated n=7, *C. difficile* infected CD4 depleted n=6. Two-way ANOVA with Tukey’s multiple comparison test. Statistical significance is indicated as follows: **** P < 0.0001.

Formation of high affinity, neutralizing antibodies is supported by CD4^+^ T follicular helper (Tfh) cells and there is limited expansion of CD4^+^ Tfh cells in the lymph nodes and spleen two weeks following C, difficile infection^24^. To determine if the antibody response to TcdA was CD4^+^ T cell dependent, mice were treated with repeated doses of an αCD4 depletion antibody or an isotype control (ISO) prior to and following infection with *C. difficile*. Mice were bled on day 35 post infection and toxin-specific serum IgG was measured. Isotype treated, *C. difficile*-infected mice generated a robust antibody response to TcdA at day 35 post-infection, while CD4-depleted mice had no detectable TcdA- or TcdB-specific serum IgG mirroring Naïve serum (**Figure 2D-F**). These data confirm that TcdA-specific antibody responses generated following infection are CD4^+^ T cell dependent and failure to generate humoral immunity against TcdB could be due to a breakdown in the TcdB-specific CD4^+^ T cell responses following infection.

### Mice generate robust CD4^+^ T cell responses following *C. difficile* infection

The quality and kinetics of the CD4^+^ T cell response following *C. difficile* infection is not well understood. Therefore, we sought to define the magnitude of the polyclonal CD4^+^ T cell response over the course of infection. There was a significant induction of CD44^hi^ CD62L^lo^ CD4^+^ T cells in the mLN of *C. difficile*-infected mice at day 21 post-infection compared to antibiotic-treated control mice (**Figure 3A,B**). (**Supplemental Figure 4C,D**). CD44 expression serves as a marker of antigen-experienced T cells and following infection the lymph node homing receptor CD62L is downregulated to enable migration to the tissue site of infection. Combined, CD44^hi^ CD62L^lo^ expression profile following infection broadly identifies activated, effector CD4^+^ T cells. Next, the mucosal CD4^+^ T cell response in the large intestine lamina propria (Li LP) were assessed. Li LP CD4^+^ T cells were primarily CD44^hi^ CD62L^lo^ expressing cells and this population of activated effector CD4^+^ T cells was significantly increased compared to antibiotic-treated control mice at days 14, 21, and 35, peaking at d14 post-infection (**Figure 3A,C**) The proportion of activated effector (CD44^hi^ CD62L^lo^) CD4^+^ T cells in the Li LP expressing Integrin α-E (CD103) increased at days 14, 21, and 35 post-infection (**Figure 3D,E**) suggesting these cells are signaling for epithelial retention and are potentially tissue-resident memory T cells^38^. Additionally, germinal Center (GC) B cells were also significantly increased at day 21 post-infection (**Supplemental Figure 4A,B**), however Tfh cells were not expanded following infection (**Supplemental Figure 4C,D**) in agreement with a previous report from Amadou Amani et al^24^. Together, these data demonstrate that polyclonal B cells and CD4^+^ T cells are being activated on a cell population level following *C. difficile* infection though the antigen specificity of these activated cells is undefined.

**Figure 3.**
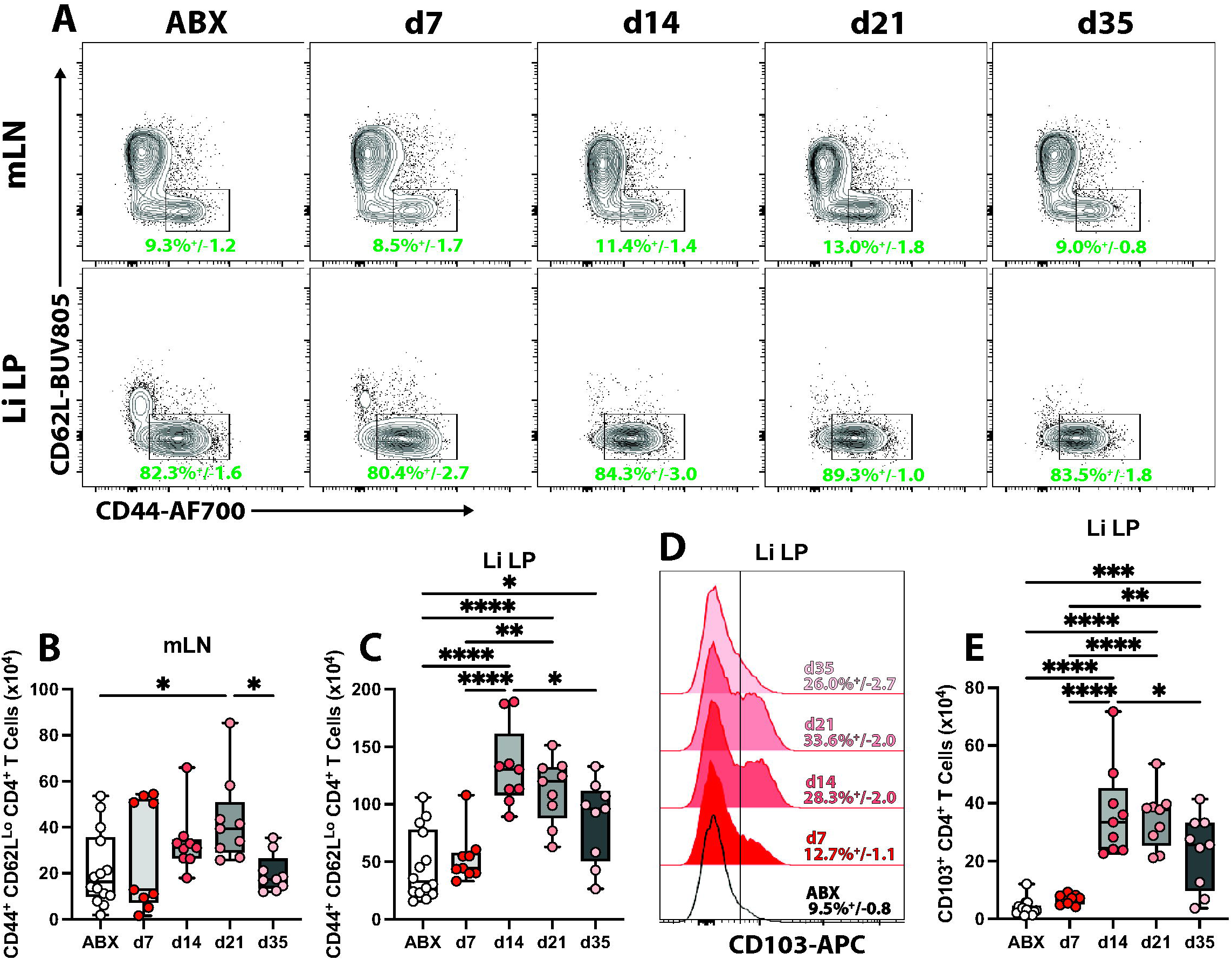
Increased activated CD4^+^ T cells following *C. difficile* infection. (A) Frequencies of activated CD44^+^ CD62L^lo^ CD4^+^ T cells from the mesenteric lymph nodes and large intestine lamina propria at various timepoints post-infection. (B) The total number of activated CD44^+^ CD62L^lo^ CD4^+^ T cells in the mesenteric lymph nodes (C) and large intestine lamina propria. (D) Frequency of activated CD4^+^ T cells expressing CD103^+^ in the lamina propria following infection. (E) Total number of CD103^+^ activated CD4^+^ T cells in the lamina propria. Representative data were pooled from two independent experiments. ABX n=14 and *C. difficile*-infected n=9 per timepoint. One-way ANOVA with Tukey’s multiple comparison test. Statistical significance is indicated as follows: * P < 0.05; ** P < 0.01; *** P < 0.001; **** P < 0.0001. Representative FACS plots were pre-gated based on the following parameters: Singlets, Live, Lymphocytes, CD45^+^, CD3/5^+^, CD4^+^. Frequency of parental gate +/- standard error of the mean is shown.

To begin to address the functionality of CD4^+^ T cells induced following infection, the cytokine profile of the Li LP CD4^+^ T cell population from *C. difficile*-infected and antibiotic-treated control mice was defined. *Ex vivo* pharmacologic stimulation with phorbol 12-myristate 13-acetate and ionomycin (PMA-I) was performed in the presence of Brefeldin A and Monensin (BFA-M) on cells isolated from the Li LP. The cytokines IFN-γ and IL-17A were assessed to measure changes to T_H_1 and T_H_17 cell populations, respectively, two major effector CD4^+^ T cell subsets in the intestine^39, 40^. IL17A^+^ CD4^+^ T cells were expanded in the Li LP of *C. difficile*-infected mice at day 7 and day 14 post-infection, with no significant induction of IFN-γ^+^ CD4^+^ T cells (**Figure 4A,B**). Thus, there is an expansion of intestinal T_H_17 cells during *C. difficile* infection, however, whether the cognate antigen of these T_H_17 cells is *C. difficile* toxin is unclear.

**Figure 4.**
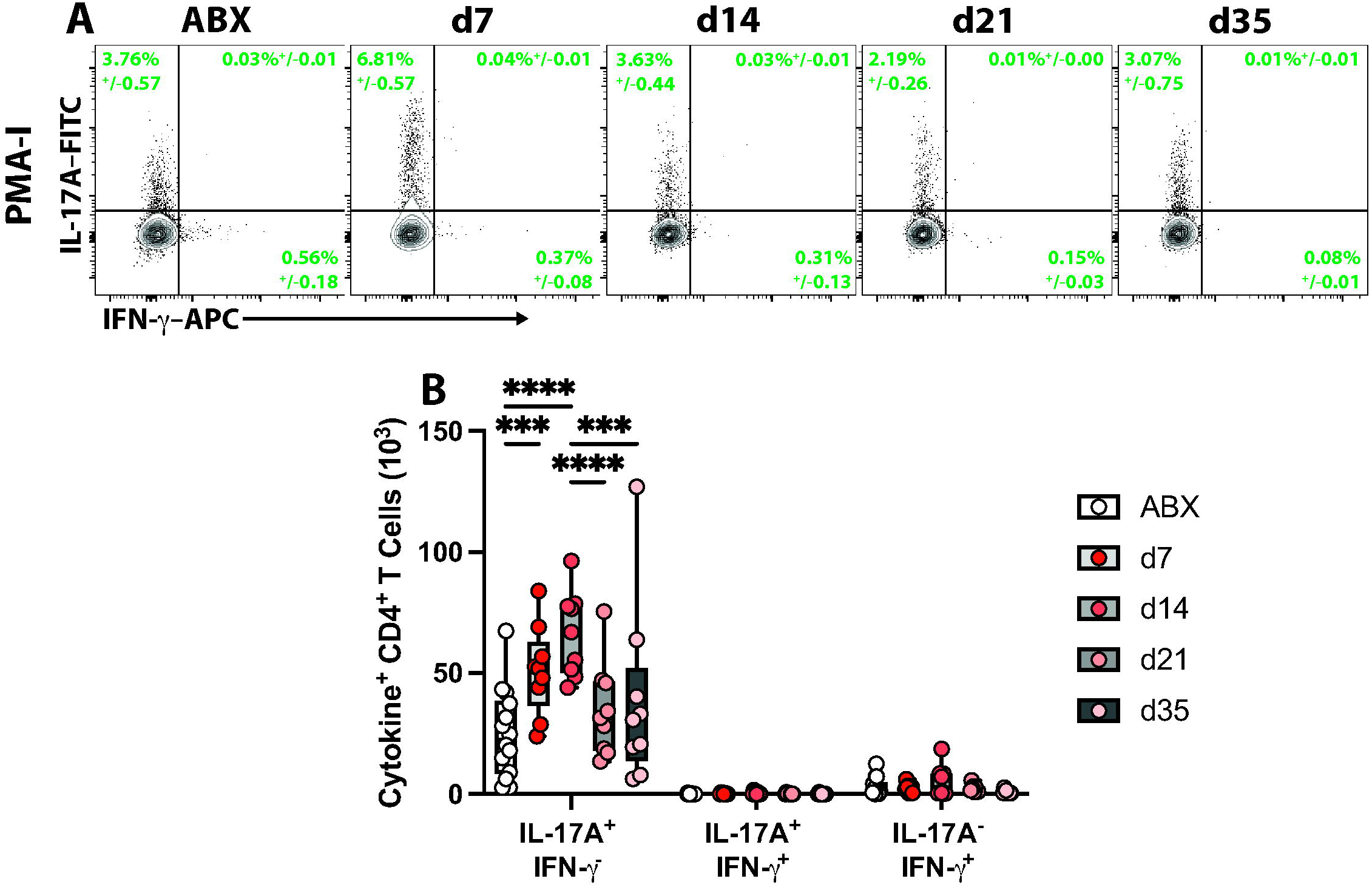
Increased T_H_17 cells in the lamina propria following *C. difficile* infection. (A) Frequencies of PMA-I stimulated CD4^+^ T cells expressing IL-17A and IFN-γ in the lamina propria. (B) Total number of cytokine-positive CD4^+^ T cells in the lamina propria following stimulation with PMA-I. Representative data were pooled from two independent experiments. ABX n=14 and *C. difficile*-infected n=9 per timepoint. Two-way ANOVA with Tukey’s multiple comparison test. Statistical significance is indicated as follows: *** P < 0.001; **** P < 0.0001. Representative FACS plots were pre-gated based on the following parameters: Singlets, Live, Lymphocytes, CD45^+^, CD3/5^+^, CD4^+^. Frequency of parental gate +/- standard error of the mean is shown.

### CD4^+^ T cells respond to TcdA but fail to respond to TcdB following infection

To determine if activated large intestinal lamina propria CD4^+^ T cells induced following *C. difficile* infection are responsive to TcdA or TcdB, peptide libraries of TcdA and TcdB were generated. Due to the large size of TcdA and TcdB (308 and 270 kDa, respectively), we sought to narrow the scope of the peptide libraries generated to relevant domains. The immune epitope database (IEDB) MHC-II binding prediction module^41, 42^ was used to assess the predicted binding of TcdA and TcdB to H2-IAb, the MHC-II molecule present in C57BL/6 mice. Upon plotting the 15mer amino acid sequence predicted binding affinity over the span of the protein, it was determined that most of the immunogenic epitopes for TcdA (**Figure 5A**) and TcdB (**Figure 5B**) were contained within the CROPs domain. Thus, 15mer peptide libraries of TcdA and TcdB CROPs domains were generated for T cell restimulation assays (**Supplemental Table 1**).

**Figure 5.**
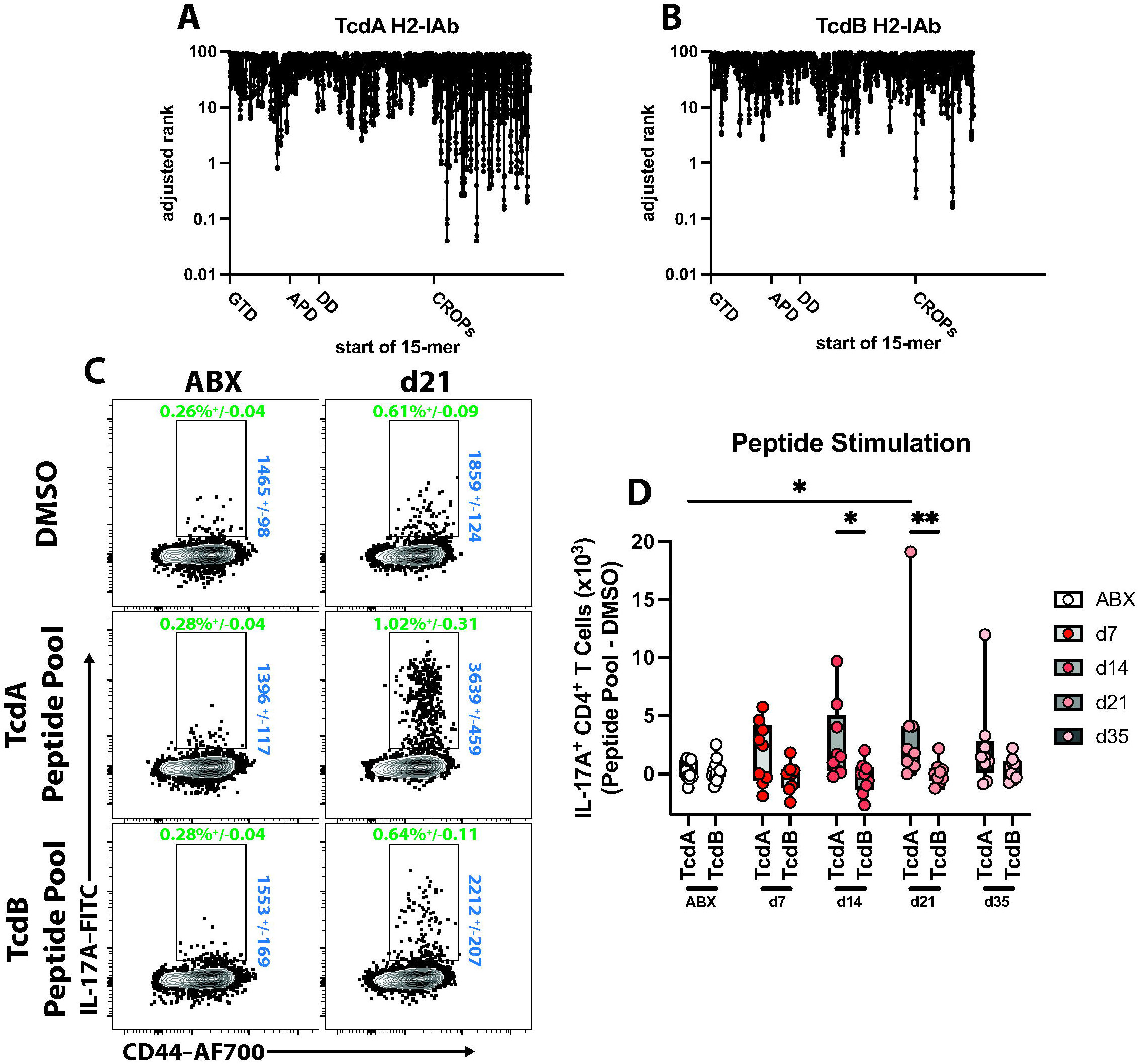
Mice generate TcdA peptide-responsive CD4^+^ T cells but not TcdB peptide-responsive CD4^+^ T cells following *C. difficile* infection. (A) Amino acid sequences from TcdA and (B) TcdB 15mers were analyzed using the Immune epitope database MHC-II peptide binding prediction for H2-IAb. Data were represented as adjusted rank which is generated by comparing each peptide’s score to a database of 15mers. Lower numbers are predicted to be better binders to H2-IAb. Peptides presented in N to C terminus order and annotated by domain. (C) Frequencies of TcdA or TcdB pooled peptide library or DMSO stimulated CD4^+^ T cells expressing IL-17A. (D) The total number of IL-17A^+^ CD4^+^ T cells responding to TcdA or TcdB peptide libraries. To account for the background cytokine production observed, the total number of IL-17A^+^ CD4^+^ T cells in the DMSO group was subtracted from the total number of peptide-responsive IL-17A^+^ CD4^+^ T cells. Representative data were pooled from two independent experiments. ABX n=14 and *C. difficile*-infected n=9 per timepoint. Two-way ANOVA with Tukey’s multiple comparison test. Statistical significance is indicated as follows: * P < 0.05; ** P < 0.01. Representative FACS plots were pre-gated based on the following parameters: Singlets, Live, Lymphocytes, CD45^+^, CD3/5^+^, CD4^+^. Frequency of parental gate +/- standard error of the mean is shown in green. IL-17A mean fluorescence intensity (MFI) +/- standard error of the mean is shown in blue. Cells were stimulated with TcdA or TcdB peptide libraries or DMSO for 1 hour before the addition of BFA-M. Cells were incubated for an additional 4 hours before intracellular cytokine staining and analysis by flow cytometry.

Cells isolated from the Li LP of antibiotic-treated or *C. difficile*-infected mice at different timepoints were restimulated with TcdA, TcdB peptide libraries, or DMSO in the presence of BFA-M. A low frequency and number of DMSO-stimulated CD4^+^ T cells produce of IL-17A that was elevated in *C. difficile* infected mice (**Supplemental Figure 5A,B**). These data establish our baseline of non-toxin specific IL-17A^+^ CD4^+^ T cell population. This baseline population is subtracted from TcdA or TcdB peptide restimulation groups to determine the number of IL-17A^+^ toxin-responsive CD4^+^ T cells. On day 21 post-infection there was a significant increase in TcdA peptide-responsive CD4^+^ T cells producing IL-17A compared to antibiotic-treated control mice (**Figure 5C,D**). In contrast, there was no significant increase in IL17A^+^, TcdB peptide-responsive CD4^+^ T cells in *C. difficile*-infected compared to antibiotic-treated control mice (TcdB: ABX vs day 21, p=0.9999) (**Figure 5C,D**). These data correspond with the humoral results, demonstrating that the lack of an antibody response to TcdB is mirrored by the absence of a CD4^+^ T cell response to this toxin.

### Glucosyltransferase activity of *C. difficile* toxins inhibits antigen-specific adaptive immune responses

TcdB is highly immunogenic, and mice generate robust TcdB-specific antibody^24–31^ and CD4^+^ T cell responses following mRNA-LNP vaccine immunization^31^. Therefore, we speculated that the active toxin produced during natural infection inhibits formation of anti-TcdB immunity. The pathology caused by *C. difficile* toxins are primarily attributed to the glucosyltransferase (GT) activity of TcdA and TcdB^9, 10, 43^. Therefore, we tested if the glucosyltransferase activity of TcdA and TcdB was inhibiting the toxin-specific adaptive immune response. To test this hypothesis, antibiotic-treated mice were infected with wild type *C. difficile* (strain R20291 WT), TcdA glucosyltransferase-mutant (TcdA_GTX_ TcdB^+^), TcdB glucosyltransferase-mutant (TcdA^+^ TcdB_GTX_), or a dual glucosyltransferase-mutant (TcdA_GTX_ TcdB_GTX_). Following infection with R20291 WT, TcdA_GTX_ TcdB^+^, TcdA^+^ TcdB_GTX_, or TcdA_GTX_ TcdB_GTX_, mice experienced disease severity similar to published reports^9^. Mice infected with TcdA_GTX_ TcdB^+^ lost slightly less weight and recovered faster compared to R20291 WT (**Figure 6A**). Mice infected with TcdA^+^ TcdB_GTX_ lost less weight compared to WT, but more weight loss compared to TcdA_GTX_ TcdB_GTX_ (**Figure 6A**). Furthermore, R20291 WT and TcdA_GTX_ TcdB^+^ infected mice experienced high mortality following infection, while no TcdA^+^ TcdB_GTX_ or TcdA_GTX_ TcdB_GTX_ infected mice succumbed to the disease (**Figure 6B**). Finally, there was no difference in bacterial burden between these infection groups, indicating that the glucosyltransferase activity of *C. difficile* toxins is not required for colonization (**Supplementary Figure 6**). These data are in agreement with previous reports and confirm that the glucosyltransferase activity of TcdB is driving lethal disease while TcdA is additive and not the primary driver of disease pathogenesis during infection^9, 10^.

**Figure 6.**
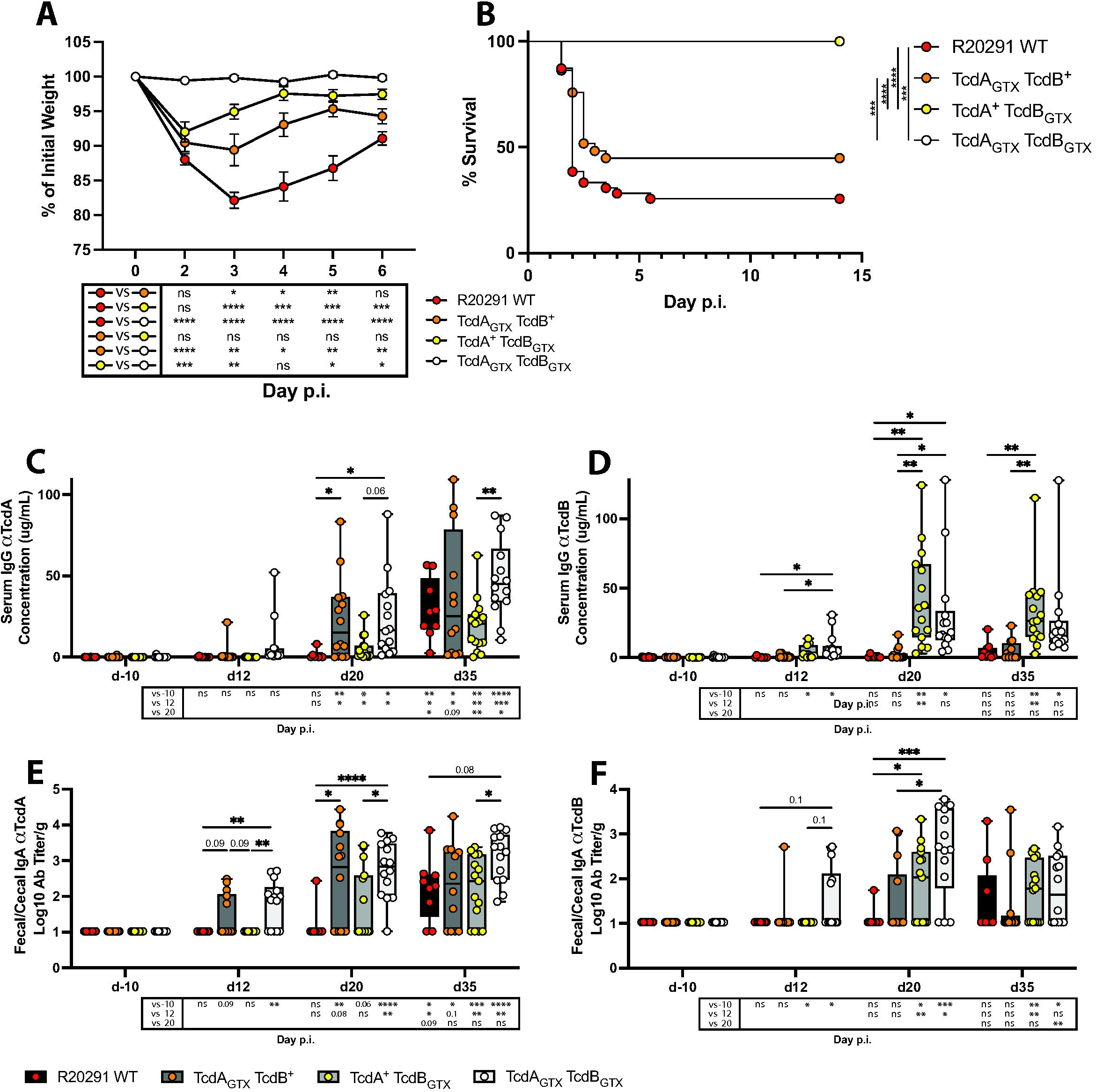
Glucosyltransferase activity of TcdA and TcdB inhibits toxin-specific serum IgG responses. (A) Weight loss and (B) Survival following infection with R20291 WT or mutant strains of *C. difficile*. (C) Quantification of serum IgG specific for TcdA-CROPs and (D) TcdB-CROPs prior to infection and following infection with *C. difficile* R20291 WT or glucosyltransferase mutants. (E) Quantification of luminal IgA specific for TcdA-CROPs or (F) TcdB-CROPs from fecal supernatants (Days -10, 12, and 20) or cecal supernatants (Day 35) prior to infection and following infection with *C. difficile* R20291 WT or glucosyltransferase mutants. Representative data were pooled from three independent experiments. WT R20291 n=9, TcdA_GTX_ TcdB^+^ n=12, TcdA^+^ TcdB_GTX_ n=15, and TcdA_GTX_ TcdB_GTX_ n=14 at day 35 post-infection. (B) Log-rank (Mantel-Cox) test with Bonferroni correction. (C-F) Two-way ANOVA with Tukey’s multiple comparison test. Statistical significance is indicated as follows: * P < 0.05; ** P < 0.01; *** P < 0.001 **** P < 0.0001.

TcdA- and TcdB-specific IgG in the serum was measured at various timepoints post-infection with R20291 WT or glucosyltransferase mutant strains. TcdA-specific IgG was induced by day 35 post-infection in mice infected with R20291 WT compared to pre-infection levels (**Figure 6C**). In contrast, there was no significant TcdB-specific IgG at day 35 post-infection with R20291 WT compared to pre-infection (**Figure 6D**) confirming that the impaired TcdB-specific antibody response was not unique to infection with the CD196 strain of *C. difficile* (**Figure 1E**).

Next, the serum TcdB-specific IgG response was compared between mice infected with the R20291 WT and three mutant strains. Supporting our hypothesis, TcdB-specific IgG in mice infected with TcdA^+^ TcdB_GTX_ and TcdA_GTX_ TcdB_GTX_ was restored compared to WT and TcdA_GTX_ TcdB^+^ infected mice at days 20 and 35 post-infection (**Figure 6D**). Infection with TcdA_GTX_ TcdB^+^ did not yield antibody titers to TcdB, mirroring the anti-TcdB IgG response in mice infected with WT R20291. Additionally, mice infected with TcdA^+^ TcdB_GTX_ and TcdA_GTX_ TcdB_GTX_ had a significantly increased TcdB-specific luminal IgA response at day 20 post infection compared to R20291 WT infected mice (**Figure 6F**). These data demonstrate that the glucosyltransferase activity of TcdB is responsible for hindering the infection-induced humoral response to TcdB.

TcdB glucosyltransferase inactivation did not alter the kinetics of the TcdA-specific antibody response (WT R20291 compared to TcdA^+^ TcdB_GTX_) (**Figure 6C,E**). However, comparison of the anti-TcdA IgG response following infection with TcdA_GTX_ TcdB^+^, TcdA_GTX_ TcdB_GTX_, and R20291 WT strains revealed an earlier induction of TcdA-specific antibodies at day 20 post-infection in mice infected with *C. difficile* strains that had the glucosyltransferase domain of TcdA inactivated (**Figure 6C**). Moreover, TcdA-specific luminal IgA was induced by day 12 post-infection in mice infected with TcdA_GTX_ TcdB_GTX_ compared to mice infected with R20291 WT and TcdA^+^ TcdB_GTX_ (**Figure 6E**). At day 20 post-infection, mice infected with either TcdA_GTX_ TcdB^+^ and TcdA_GTX_ TcdB_GTX_ had TcdA-specific IgA titers that were significantly greater than the R20291 WT group (**Figure 6E**). Together, these data suggest that the glucosyltransferase activity of TcdA is delaying the systemic IgG and mucosal IgA response to TcdA but there is no significant cross-toxin inhibition from TcdB glucosyltransferase activity on the TcdA-specific humoral response. Moreover, the TcdA_GTX_ TcdB_GTX_ infection group exhibited an earlier and significant induction of both TcdA and TcdB IgG and IgA responses compared to R20291 WT IgG and IgA responses (**Figure 6C-F**), confirming that ablation of both TcdA and TcdB glucosyltransferase activity is required to restore both toxin-specific antibody responses.

To determine if toxin-specific CD4^+^ T cell responses were analogously inhibited by GT activity of TcdA and TcdB, *ex vivo* TcdA, or TcdB peptide library restimulation were performed on cells isolated from the lamina propria of mice on day 14 post infection with R20291 WT, TcdA_GTX_ TcdB_GTX_, or antibiotic-treated control mice. Following TcdA peptide library stimulation, mice infected with TcdA_GTX_ TcdB_GTX_ exhibited an increase in IL-17A+ CD4+ T cells compared to antibiotic-treated control mice (**Figure 7A**). Further there was a marked expansion of TcdA peptide-responsive CD4^+^ T cells that produced IFN-γ and TNF-α compared to antibiotic-treated control mice or R20291 WT-infected mice (**Figure 7B**). More TcdA-responsive CD4^+^ T cells isolated from TcdA_GTX_ TcdB_GTX_ infected mice were polyfunctional, dual cytokine producing cells compared to TcdA-responsive CD4^+^ T cells isolated from R20291 WT infected mice (**Figure 7C**). Following TcdB peptide library stimulation, mice infected with TcdA_GTX_ TcdB_GTX_ saw a significant induction of IL-17A^+^, IFN-γ^+^, and TNF-α^+^ CD4^+^ T cells compared to antibiotic-treated control mice and R20291 WT-infected mice (**Figure 7A,B,D**). Therefore, TcdA and TcdB are inhibiting the toxin-specific CD4^+^ T cell response. This mechanism is dependent on the glucosyltransferase activity of these toxins, as infection with a glucosyltransferase mutant strain restores the toxin-specific CD4^+^ T cell response.

**Figure 7.**
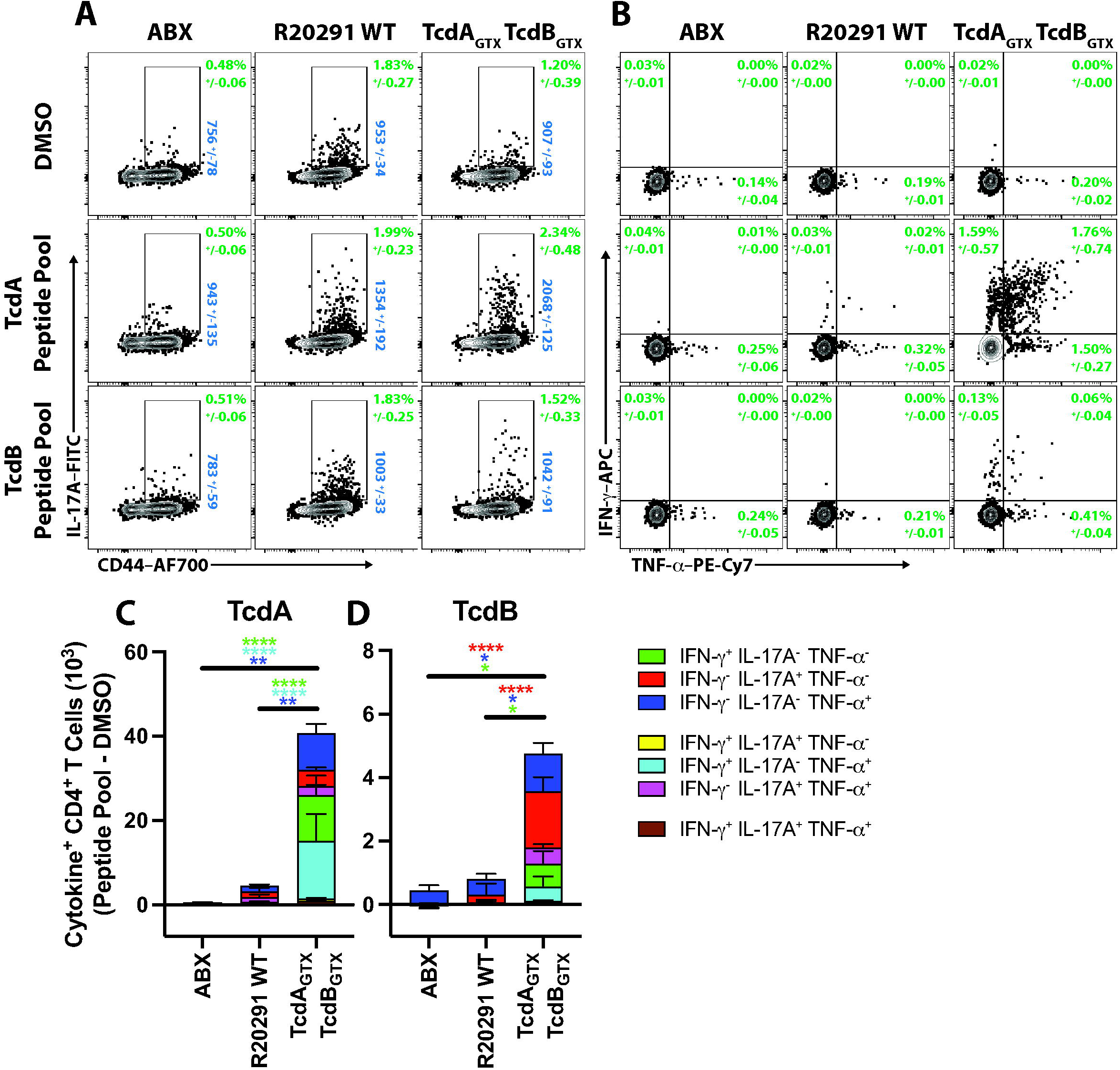
Glucosyltransferase activity of TcdA and TcdB inhibits the toxin-responsive CD4^+^ T cell response. (A) Frequencies of TcdA or TcdB peptide pool-stimulated or DMSO-stimulated CD4^+^ T cells isolated from the LiLP of antibiotic-treated control mice, or mice infected with R20291 WT or TcdA_GTX_ TcdB_GTX_ expressing IL-17A or (B) IFN-γ and TNF-α on day 14 post infection. (C) The total number of cytokine^+^ CD4^+^ T cells responding to TcdA or (D) TcdB peptide libraries. To account for the background cytokine production observed, the total number of cytokine^+^ CD4^+^ T cells in the DMSO group was subtracted from the total number of peptide-responsive CD4^+^ T cells. ABX n=5, R20291 WT infected n=9, and TcdA_GTX_ TcdB_GTX_ infected n=5. Two-way ANOVA with Tukey’s multiple comparison test. Statistical significance is indicated as follows: ** P < 0.01; *** P < 0.001; **** P < 0.0001. Representative FACS plots were pre-gated based on the following parameters: Singlets, Live, Lymphocytes, CD45^+^, CD3/5^+^, CD4^+^. Frequency of parental gate +/- standard error of the mean is shown in green. IL-17A mean fluorescence intensity (MFI) +/- standard error of the mean is shown in blue. Cells were stimulated with TcdA or TcdB peptide libraries or DMSO for 1 hour before the addition of BFA-M. Cells were incubated for an additional 4 hours before intracellular cytokine staining and analysis by flow cytometry.

## Discussion

This report defines the humoral and cellular adaptive immune responses to both TcdA and TcdB following *C. difficile* infection. Infected mice generated robust antibody responses to TcdA in the serum that peaked 35 days following infection but failed to respond to TcdB. These data were mirrored in the intestine with luminal IgA where mice generated high titers of TcdA-specific IgA with minimal TcdB-specific IgA. These data are in agreement with previous studies that suggested a lack of TcdB-specific antibodies following primary infection were associated with susceptibility in relapse infection models^24, 25^. Other groups have demonstrated that mice generate antibody responses to TcdA following infection^44, 45^; however, this report is the first to combine these findings, showing that the TcdA-specific adaptive immune response is largely intact while there is a failed TcdB-specific humoral immune response following *C. difficile* infection.

Activated CD4^+^ T cells were expanded in the large intestine lamina propria with a significant increase in T_H_17 cells following *C. difficile* infection. Further, TcdA-responsive, IL-17A-producing CD4^+^ T cells were significantly induced compared to antibiotic-treated control mice, but TcdB-responsive CD4^+^ T cells were undetected. These data confirm that the failure to generate TcdB-specific antibody responses is paralleled by a failure to induce TcdB-specific CD4^+^ T cells. This report is the first to identify toxin-responsive CD4^+^ T cells in the intestine following *C. difficile* infection. Further work is required to determine if these T_H_17 skewed responses following infection are protective or pathogenic. IL-17A is a pluripotent cytokine during *C. difficile* infection. T_H_17 cells from DSS-treated mice adoptively transferred prior to *C. difficile* infection significantly increased severity from subsequent infection suggesting that T_H_17s are disadvantageous^46^. However, IL-17A appears to be protective during acute *C. difficile* infection as neutralizing or genetically ablating IL-17A increases disease severity and reduces survival^47^.

To determine the toxin-intrinsic mechanism for failed toxin-specific adaptive immune responses, *C. difficile* mutants with ablated glucosyltransferase activity of TcdA and/or TcdB were used^9^. These mutants reveal that the glucosyltransferase activity of TcdA and TcdB inhibit both the humoral and cellular toxin-specific immune response to TcdA and TcdB, respectively. This report is the first to demonstrate immunomodulatory effects of the glucosyltransferase activity of *C. difficile* toxins and their effects on the toxin-specific adaptive immune response.

There are multiple cellular processes that could be targeted by the glucosyltransferase activity of the toxin that could result in the inhibition of adaptive immune responses. TcdA and TcdB drive cell death of cultured monocytes, macrophages, and T cells^48–55^. Therefore, immune cell apoptosis from TcdA and TcdB may dampen the adaptive immune response to these toxins. While similar in structure and function, TcdB is the primary virulence factor for *C. difficile*^12^ and is 1000x more cytotoxic than TcdA^56^. Therefore, the increased cytotoxicity of TcdB may drive cell death of immune cells *in vivo*, which could lead to the reduced antibody response to TcdB over TcdA. Alternatively, the mechanism of action could be through inhibition of cellular processes essential for successful adaptive immune responses. Notably, the cellular targets for TcdA and TcdB are host GTPase enzymes such as Rho, which regulate the assembly and organization of actin^6, 7, 57^. Inhibition of host GTPase enzymes by *C. difficile* toxins leads to a loss of cellular morphology due to the breakdown of the actin cytoskeleton. These processes are critical for cellular migration^7^. Inhibition of host GTPase enzymes in an intoxicated antigen-presenting cell might prevent it from migrating to a secondary lymphoid organ, subsequently preventing it from presenting toxin antigen to cognate T cells. Furthermore, inhibition of GTPases by treating spleen-derived dendritic cells or bone-marrow derived dendritic cells with TcdB completely blocked the *in vitro* macropinocytic antigen uptake^58, 59^ as well as endocytic activity in dendritic cells^59, 60^. Finally, TcdB treatment of bone-marrow derived dendritic cells has been demonstrated to hinder antigen presentation to cognate T cells *in vitro*^60^. Therefore, the targeted inhibition of GTPases by TcdA and TcdB in dendritic cells could directly prevent subsequent antigen uptake, processing, migration, and presentation of the toxins to cognate CD4^+^ T cells. Alternatively, a recent report by Norman *et al.* determined that TcdB inhibits memory B cell recall responses following vaccination through CXCR4 upregulation on B cells^61^ suggesting glucosyltransferase-mediated inhibition of humoral immunity following infection could be directly attributed to B cell migration.

The culmination of human correlative studies, anti-toxin monoclonal antibody treatment, and murine vaccine models have demonstrated that antibodies to *C. difficile* toxins effectively prevent severe primary and recurrent disease. However, it is unclear why some patients fail to generate adaptive immune responses, leaving them susceptible to symptomatic disease recurrence. The results from this study strongly support the notion that patients fail to generate antibodies due to the glucosyltransferase activity of TcdA and TcdB. Furthermore, our data suggest patients that generate antibody titers are likely still immunologically hindered by toxin glucosyltransferase activity. A study by Shah *et al.*, observed that mAbs generated from TcdB-specific B cells isolated from patients have low to medium affinity to TcdB and are poor neutralizers^62^. This suggests that even when patients generate antibody responses to TcdB, antibodies elicited are poorly neutralizing with low somatic mutations. Inhibiting the glucosyltransferase activity of TcdA and TcdB during *C. difficile* infection holds potential for unleashing the adaptive immune response to facilitate recognition, and sustained memory responses to *C. difficile* toxins, and ultimately prevent symptomatic disease recurrence.

## Methods

### Mice

C57BL/6 mice were bred and maintained at the University of Pennsylvania under specific pathogen-free conditions. Mice were maintained in autoclaved cages with autoclaved food (LabDiet 5010) and water ad libitum. Sex and age-matched controls were used in all experiments according to institutional guidelines for animal care. All animal procedures were approved by the Institutional Animal Care and Use Committee of the University of Pennsylvania prior to initiation of studies (Protocol Number: 806371).

### Antibiotic pretreatment and *C. difficile* infection

Two- to four-month-old mice were infected with *C. difficile* spores using a previously described infection protocol^32, 33^. Briefly, seven days before infection, mice were administered mice were administered neomycin (0.33g/L), metronidazole (0.25g/L), and vancomycin (0.33g/L) ad libitum in the drinking water for four days then given sterile water for the remainder of the experiment. One day before infection, mice were administered 200µg of clindamycin by intraperitoneal injection. Twenty-four hours later, mice received 1000 *C. difficile* spores (CD196 or WT or mutant R20291 strains) via oral gavage. After infection, mice were monitored daily for disease by weight loss and morbidity scoring. Mice were humanely euthanized if they had signs of fatal disease including lethargy, hunched posture, were moribund, or weight loss of more than 30% of the pre-infection weight.

### Quantification of *C. difficile* burden and toxin titers

Fecal pellets or cecal content were resuspended in deoxygenated PBS in an anaerobic chamber (Coy Labs). Ten-fold serial dilutions in deoxygenated PBS were plated on Difco Brain Heart Infusion (BD Biosciences) agar plates supplemented with 5 g/L yeast extract (BD Biosciences), 1 g/L L-cysteine (Sigma), 4 g/L D-cycloserine (Sigma), 16 µg/L cefoxitin (Sigma), and 1 g/L taurocholate (MP Biomedicals) overnight at 37°C under anaerobic conditions^63^. Counts are represented as Log10 and normalized to the weight of the starting material. Resuspended fecal and cecal samples were centrifuged at 13,000g for 5 minutes, and supernatants were used to quantify *C. difficile* toxin titers. Vero cells (monkey kidney epithelial cells [ATCC# CCL-81]) plated on flat bottom tissue culture treated 96 well plates (Corning) at 2x10^5^ cells/mL in complete medium (DMEM [Corning] supplemented with 10% FBS, 1% penicillin/streptomycin [100 U/mL and 100 µg/mL, respectively] [Gibco], 50 µg/mL gentamicin [Gibco], 10 mM HEPES [Gibco], 0.5 mM β-mercaptoethanol [Gibco], 1mM Sodium Pyruvate [Sigma], 20 µg/mL L-glutamine [Corning]). Cells were incubated at 37°C overnight in a humidified atmosphere at 5% CO_2_. Next, serial dilutions of cecal supernatant diluted in PBS were added to the Vero cell monolayers and incubated at 37°C for 24 hours. To ensure cell rounding was due to *C. difficile* toxins, samples were also incubated with antitoxin antisera (Techlab) at 37°C for 1 hour before being transferred to Vero cell monolayers. The next day, cells were observed by light microscopy for the presence of cell rounding. The highest dilution where cell rounding occurred was designated as the toxin titer. Samples where no cell rounding was observed were set as 1/4^th^ the limit of detection. The data are represented as Log10 and normalized to the weight of the starting material.

### Generation of recombinant TcdA and TcdB proteins

The generation of TcdA-CROPs and TcdB-CROPs proteins was performed as previously described^31^. In brief, codon-optimized TcdA-CROPs and TcdB-CROPs sequences were synthesized by Genscript and cloned into the pET30a vector. An N-terminal 6XHIS tag followed by the TEV recognition/cleavage site was introduced after the start codon. Positive cultures were grown, and supernatants were concentrated and purified using a NI-AID column. Proteins were cleaved with TEV protease, buffer exchanged into PBS with 5% sucrose, quantified using micro-BCA, and the purity was determined using densitometric analysis of a Coomassie blue stained SDS-PAGE gel under reducing conditions. Both proteins displayed more than 90% purity and had the predicted molecular weights using Western blot.

### Anti-toxin antibody ELISAs

Antibodies to TcdA and TcdB were assessed by indirect ELISA. High-binding polystyrene plates (Corning) were coated overnight at 4°C with 100uL of recombinant TcdA-CROP, TcdB-CROP, full-length TcdA (The Native Antigen Company), or full-length TcdB (Abcam) proteins at 0.5µg/mL in coating buffer (sodium bicarbonate [2.93 g/L; Sigma] and sodium carbonate [1.59 g/L; Sigma] in dH_2_O at pH:9.6.). Plates were washed three times with wash buffer (1x PBS with Tween-20 [0.05%v/v; Sigma]). Plates were blocked at room temperature in 200µL of blocking buffer (1x PBS with bovine serum albumin [2%w/v; Sigma]) for 2 hours. Plates were washed three times with wash buffer before adding of 100µL of serially diluted samples diluted in blocking buffer. Pre-weighted fecal or cecal samples were resuspended in 1mL of PBS, and centrifuged at 10,000g for 5 minutes to pellet debris before dilutions. Plates were incubated overnight at 4°C with samples and then washed three times before the addition of a secondary antibody. For IgG, 100µL of Horseradish Peroxidase conjugated Donkey Anti-Mouse IgG (H+L) (1:10,000 v/v in blocking buffer; Jackson ImmunoResearch) was incubated at room temperature for 1 hour. For detection of IgA, 100µL of Biotin conjugated Rat Anti-Mouse IgA (1:3000v/v in blocking buffer; BioLegend: clone RMA-I) was added for 1 hour, washed 3 times, then incubated with streptavidin-HRP (1:10,000 v/v in blocking buffer; BD Biosciences) for 1 hour. Samples were washed three times in wash buffer before adding TMB Substrate (BD Biosciences). Plates were developed for 10 minutes before adding 2N H_2_SO_4_ (Sigma) to quench the reaction. Absorbance at 450nm was measured on SpectraMax 190 plate reader (Molecular Devices). To quantify the amount of toxin-specific IgG in serum, monoclonal antibodies to TcdA (Invitrogen: clone A73H) or TcdB (Invitrogen: clone A13I) were used to generate standard curves, and optical densities were compared back to the standard curve to quantify the concentration of toxin-specific IgG present. To quantify the amount of toxin-specific cecal IgA, end-point dilution titer was used and defined as the highest dilution of cecal supernatant that resulted in an OD greater than the cut-off OD value determined using the Frey Method^64^ with a significance level of 0.0001%. Following enumeration, the fecal antibody titers were normalized to the gram of content collected. IgA samples that were below the limit of detection were set at 1/4^th^ the average limit of detection.

### Toxin neutralization assay

Vero cells were plated on flat bottom tissue culture treated 96 well plates (Corning) at 2x10^5^ cells/mL in complete medium. Cells were incubated at 37°C overnight in a humidified atmosphere at 5% CO_2_. Next, serial dilutions of serum diluted in complete media were added to a round bottom plate (Corning) with 10pM TcdA (The Native Antigen Company) or 2pM TcdB (Abcam) and incubated at 37°C for 1 hour. As a positive control, TcdA or TcdB was incubated with 1µg/mL Actoxumab or Bezlotoxumab (MedChemExpress) to confirm neutralization capacity. Next, Vero cell supernatants were aspirated, and the serum toxin mixture was added to the Vero cell monolayers. The cells were incubated at 37°C overnight. The next day, cells were assessed for the presence of cell rounding. The neutralization titer is expressed as the last dilution of serum where protection from cell rounding occurred. Samples where no protection was observed were set at 1/4^th^ the limit of detection. Representative brightfield images were taken with a 20x air objective on a confocal microscope (Nikon) and analyzed using ImageJ version 1.54f.

### CD4^+^ T cell depletion

*In vivo* CD4^+^ T cell depletion was modified from a previously described protocol^65^. Briefly, 500 μg of the αCD4 monoclonal antibody (BioXcell, GK1.5) or an isotype control antibody (BioXcell, TNP687) was administered by intraperitoneal injection on day -1 and day 0 following the standard *C. difficile* infection protocol and readministered every 12 days to maintain CD4^+^ T cell depletion.

### Isolation of immune cells from lymph nodes and lamina propria

Single cell suspensions were obtained from the mesenteric lymph nodes by mechanical separation. Lymph nodes were dissected, placed on a 100um cell strainer, and crushed with a 3mL syringe to dissociate the tissue. To isolate single-cell suspensions from the large intestine lamina propria (Li LP), the large intestine and caecum were dissected, cut longitudinally, and washed with PBS. To remove epithelial cells, epithelial strip buffer (PBS, 5mM EDTA, 1mM dithiothreitol, 4% FBS, 1% penicillin/streptomycin) was added to the tissue before incubation at 37°C while shaking at 180 rpm for 10 minutes. Incubation in additional epithelial strip buffer for 20 minutes removes intraepithelial lymphocytes. The remaining tissue was digested in collagenase IV (1.5 mg/mL [500U/mL]), DNase (20 µg/mL) in complete tissue culture media (RMPI supplemented with 10% FBS, 1% penicillin/streptomycin, 50 µg/mL gentamicin, 10 mM HEPES, 0.5 mM β-mercaptoethanol, 1mM Sodium Pyruvate, 20 µg/mL L-glutamine) at 37°C while shaking at 180 rpm for 30 minutes. Supernatants containing the Li LP fraction were passed through a 100µm cell strainer, pelleted by centrifugation at 500g, resuspended, and passed through a 40µm cell strainer. Cells were centrifuged at 500g and resuspended in complete media for subsequent analysis.

### Generation of Peptide libraries

TcdA and TcdB were analyzed for predicted binding affinity for the C57BL/6 MHC-II, H2-IAb. We utilized the immune epitope database (IEDB) MHC-II binding module^41, 42^. The amino acid sequence of TcdA and TcdB from CD196^66^ 15mers were assessed for predicted binding affinity for H2-IAb. To generate peptide libraries of TcdA and TcdB, 15mer (11 amino acid overlap) peptide libraries were generated from TcdA (1831-2710) and TcdB (1835-2366) by Genscript.

### Cell stimulation and flow cytometry

To detect cytokine production from CD4^+^ T cells, cells in a single cell suspension were cultured in a flat bottom non-tissue culture treated 96 well plate (Corning) in complete medium. Cells were cultured for 1 hour at 37°C in the presence of TcdA peptide pools, TcdB peptide pools (2.5µg/mL/peptide), with DMSO as a negative control, or PMA (0.05 ug/mL [Sigma]) and Ionomycin (0.5 ug/mL [Sigma]) as a positive control. After 1 hour, Brefeldin-A and Monensin (eBioscience) were added and incubated for an additional 4 hours to allow for cytokine accumulation. Following incubation (or directly from single-cell suspension for surface staining), cells were transferred to 96 well round bottom plates for cell staining for flow cytometry using a standard protocol. Briefly, cells were washed in PBS, stained with Zombie Aqua (BioLegend) at room temperature for 10 minutes, and washed in FACS buffer (PBS, 1% BSA, 50µg/mL Sodium Azide) before blocking at 4°C with anti-CD16/32 antibody (BD Biosciences) and rat IgG (Sigma) for 20 minutes. Next, antibodies against surface antigens were added and stained for 30 minutes at 4°C. Surface antibodies included CD4−BUV395 (BD Biosciences, RM4-5), CD62L−BUV805 (BD Biosciences, MEL-14), CD45−BV605 (BioLegend, 30-F11), CD19−BV650 (BioLegend, 6D5), Ly6G−BV785 (BioLegend, 1A8), CD3−PerCP-Cy5.5 (eBioscience, 1452C11), CD5−PerCP-Cy5.5 (eBioscience, 53-7.3), CD44−AF700 (BioLegend, IM7), PD-1−PE-Cy7 (eBioscience, J43). FAS−BV605, (BioLegend, SA367H8), GL7−FITC (BioLegend, GL7). GR-1−AF700 (BioLegend, RB6-8C5). For Tfh staining, cells were incubated with a biotinylated CXCR5 antibody (eBioscience, SPRCL5) for 45 minutes at 4°C, and washed before the addition of fluorophore conjugated streptavidin (Streptavidin−BV421 [BioLegend]) with all other surface antibodies. Following surface staining, cells were fixed using 2% paraformaldehyde solution for surface staining only or intracellular cytokine fixation buffer for intracellular staining (eBioscience) for 30 minutes at 4°C. Intracellular antibodies were diluted in 1x permeabilization buffer (eBioscience) and stained intracellular antigens for 30 minutes at 4°C. Intracellular antibodies included IL17A−FITC (BioLegend, TC11-18H10.1), IFN-γ−APC (eBioscience, XMG1.2), and TNF-α−PE-Cy7 (MP6-XT22, eBioscience). Cells were washed and then analyzed on a BD Symphony A5 flow cytometer or a BD Symphony A3 flow cytometer. Flow cytometry data analysis was performed using FlowJo version 10.10.

### Data analysis

Data was analyzed using Microsoft Excel version 16.81 and transferred into GraphPad Prism version 10.0.0 for data visualization and statistical analysis. One-way ANOVA, two-way ANOVA, or mixed-effects models were utilized depending on the independent variables present and are indicated in figure legends. Before pooling antibiotic-treated control samples, a one-way ANOVA was performed and found no statistical difference between timepoints for any dataset presented. Statistical significance is indicated as follows: * P < 0.05; ** P < 0.01; *** P < 0.001; **** P < 0.0001. Figures were generated in Adobe Illustrator version 27.7. The infection model was made using BioRender.

## Author contributions

J.R.M. and M.C.A. contributed to experimental design. J.R.M., J.A.L., J.E.D., E.N.H., N.V.D., and M.Z.A. performed the experiments. F.C.P.G., M.G.A., D.B.L., and J.P.Z. assisted with reagents, shared *C. difficile* strains, and protocols. J.R.M. analyzed experimental data and generated figures. J.R.M. and M.C.A. wrote the manuscript. All authors have read, discussed, and edited the manuscript and approved the submitted version.

## Acknowledgements

The authors thank members of the Abt Lab for their critical review of the manuscript and thoughtful discussions of the results. We thank Garima Dwivedi at the Penn Institute for RNA Innovation for assisting with peptide library acquisition and sharing of protocols.

## Data availability statement

All data generated or analyzed during this study are publicly available from the corresponding author on upon publication.

## Supplemental Figure Legends

**Supplemental Figure 1. *C. difficile* infection model.** (A) Mice were infected with *C. difficile* spores (CD196 strain) following antibiotic pretreatment. (B) *C. difficile* burden, (C) and toxin titers in the cecal content following *C. difficile* infection. Representative data were pooled from two independent experiments. ABX n=14 and *C. difficile*-infected n=9 per timepoint. One-way ANOVA with Tukey’s multiple comparison test. Statistical significance is indicated as follows: **** P < 0.0001.

**Supplementary Figure 2. Lack of serum antibody repose to full-length TcdB.** (A) The optical density of serum IgG specific for TcdA or (B) TcdB at various timepoints post *C. difficile* infection. (C) Quantification of serum IgG specific for TcdA and TcdB. Representative data were pooled from two independent experiments. ABX n=14 and *C. difficile*-infected n=9 per timepoint. Two-way ANOVA with Tukey’s multiple comparison test. Statistical significance is indicated as follows: **** P < 0.0001.

**Supplementary Figure 3. Actoxumab and Bezlotoxumab demonstrate neutralization capacity *in vitro*.** Representative brightfield images of Vero cell monolayers demonstrating neutralization capacity of TcdA (10pM) and TcdB (2pM) with Actoxumab and Bezlotoxumab, respectively.

**Supplementary Figure 4. CD4^+^ T cell expansion following *C. difficile* infection.** (A) Frequency and (B) total number of GC B cells and (C,D) Tfh cells from the mesenteric lymph nodes at various timepoints post-infection. Data were pooled from two independent experiments. ABX n=14 and *C. difficile*-infected n=9 per timepoint. One-way ANOVA with Tukey’s multiple comparison test. Statistical significance is indicated as follows: * P < 0.05; ** P < 0.01; *** P < 0.001; **** P < 0.0001. Representative Tfh cell flow plots were pre-gated based on the following parameters: Singlets, Live, Lymphocytes, CD45^+^, CD3/5^+^, CD4^+^, CD44^+^ CD62L^-^. Representative GC B cell flow plots were pre-gated based on the following parameters: Singlets, Live, Lymphocytes, CD45^+^, GR1^-^, CD3^-^, CD19^+^. Frequency of parental gate +/- standard error of the mean is shown.

**Supplementary Figure 5. Increased baseline IL-17A^+^ CD4^+^ T cells following *C. difficile* infection.** (A) Frequency and (B) total number of DMSO stimulated IL-17A^+^ CD4^+^ T cells expressing IL-17A in the large intestine lamina propria following *C. difficile* infection. ABX n=5, R20291 WT infected n=9, and TcdA_GTX_ TcdB_GTX_ infected n=5. Two-way ANOVA with Dunnett’s multiple comparisons test. Statistical significance is indicated as follows: * P < 0.05; ** P < 0.01. Representative FACS plots were pre-gated based on the following parameters: Singlets, Live, Lymphocytes, CD45^+^, CD3/5^+^, CD4^+^. Frequency of parental gate +/- standard error of the mean is shown.

**Supplementary Figure 6. Bacterial burden is independent on TcdB glucosyltransferase activity.** *C. difficile* burden following infection with the parental R20291 WT strain or glucosyltransferase mutants. Data were pooled from four independent experiments. R20291 WT: n=39, TcdA_GTX_ TcdB^+^: n=30, TcdA^+^ TcdB_GTX_: n=20, TcdA_GTX_TcdB_GTX_: n=19. Mixed effects analysis with Tukey’s multiple comparison test. Statistical significance is indicated as follows: * P < 0.05; ** P < 0.01; *** P < 0.001; **** P < 0.0001.

**Supplementary Figure 7. Gating strategy for CD4^+^ T cells and GC B cells.** (A) Representative gating strategy shown for CD4^+^ T cells. Lamina propria cells from a naïve mouse are shown. (B) Representative gating strategy for GC B cells. mLN cells from a naïve mouse are shown.

**Supplementary Table 1. Peptide library.** (A) TcdA-CROP and (B) TcdB-CROP peptide libraries are shown. Peptide libraries are 15mers with 11 amino acid overlapping regions starting from amino acid numbers 1832 and 1836 for TcdA and TcdB, respectively.

