## Supplemental Figures & Table for "*Clostridioides difficile* toxin A and toxin B inhibit toxin-specific adaptive immune responses through glucosyltransferase-dependent activity"

**Supplemental Figure 1**

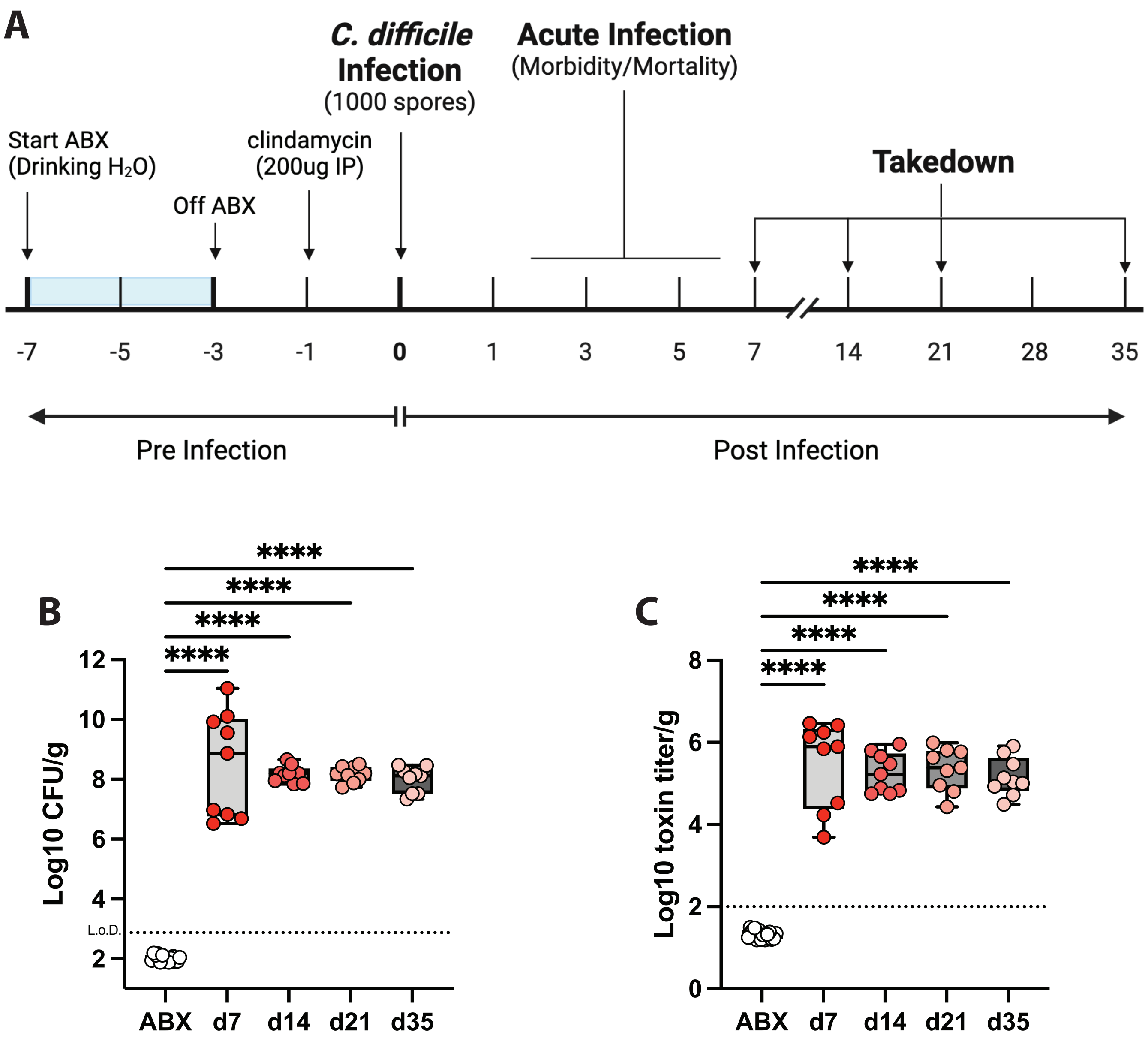

**Supplemental Figure 2**

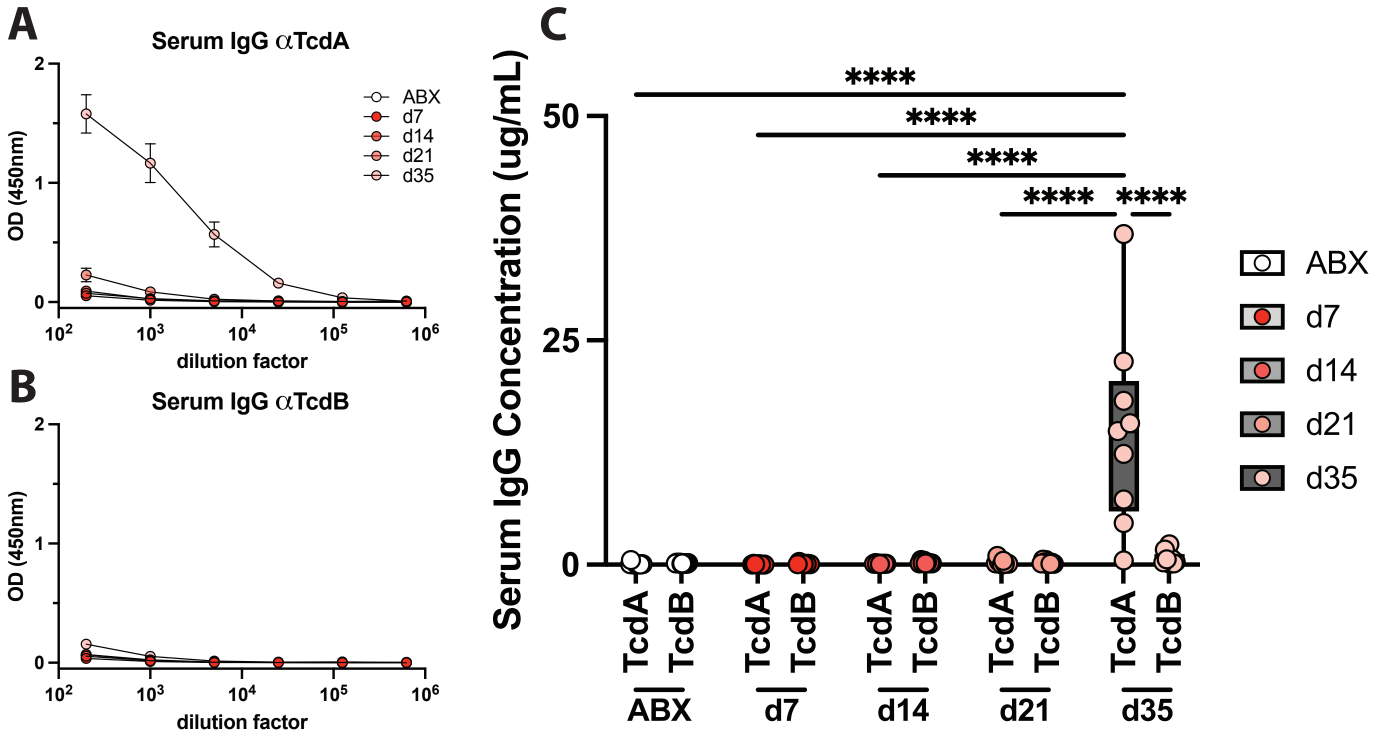

**Supplemental Figure 3**

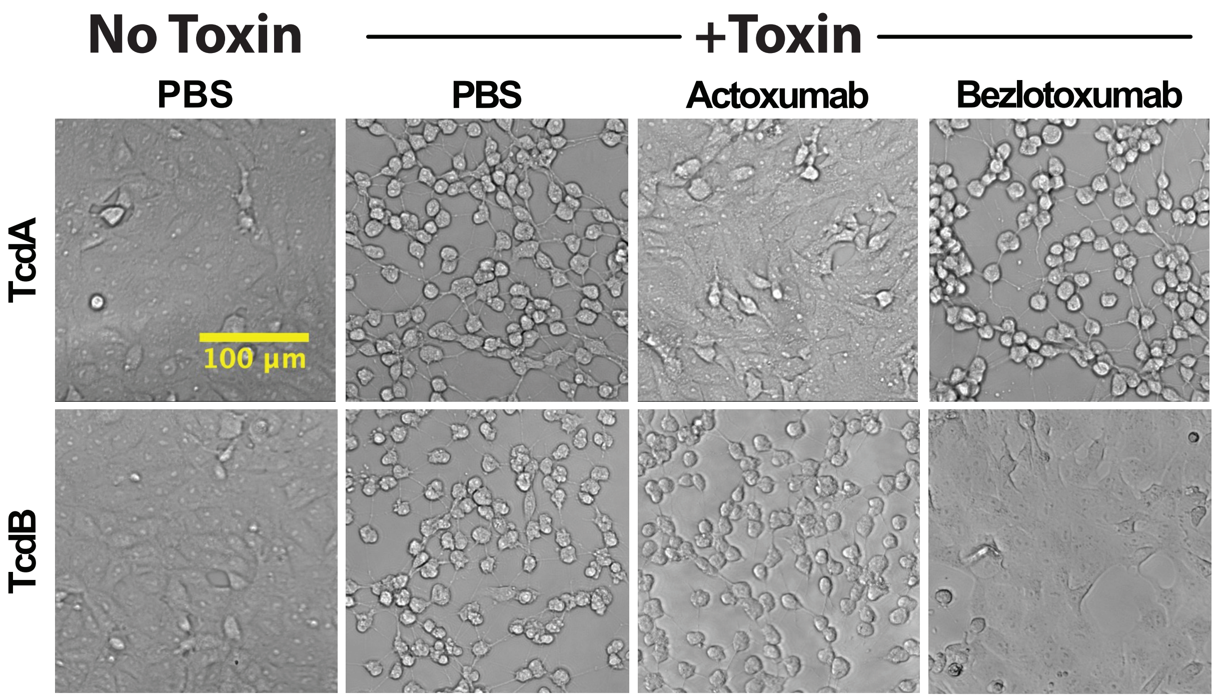

**Supplemental Figure 4**

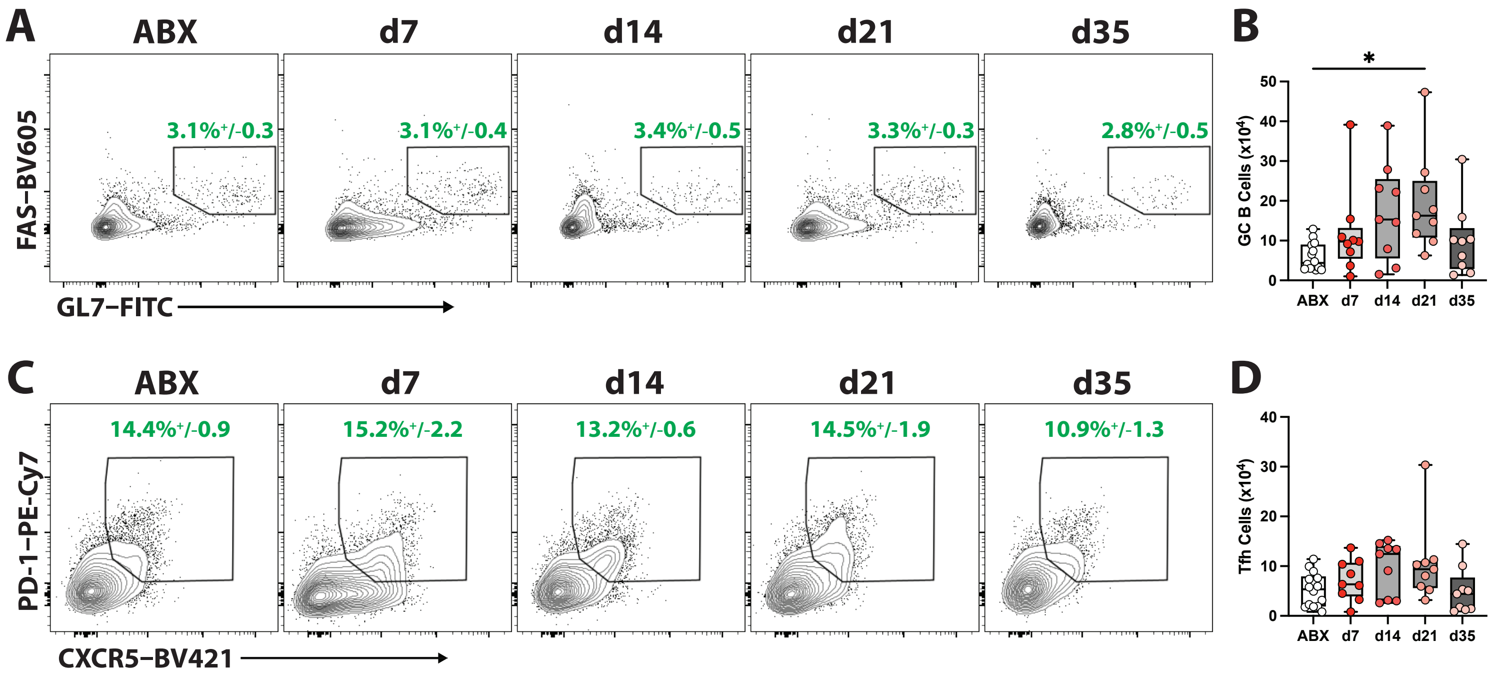

**Supplemental Figure 5**

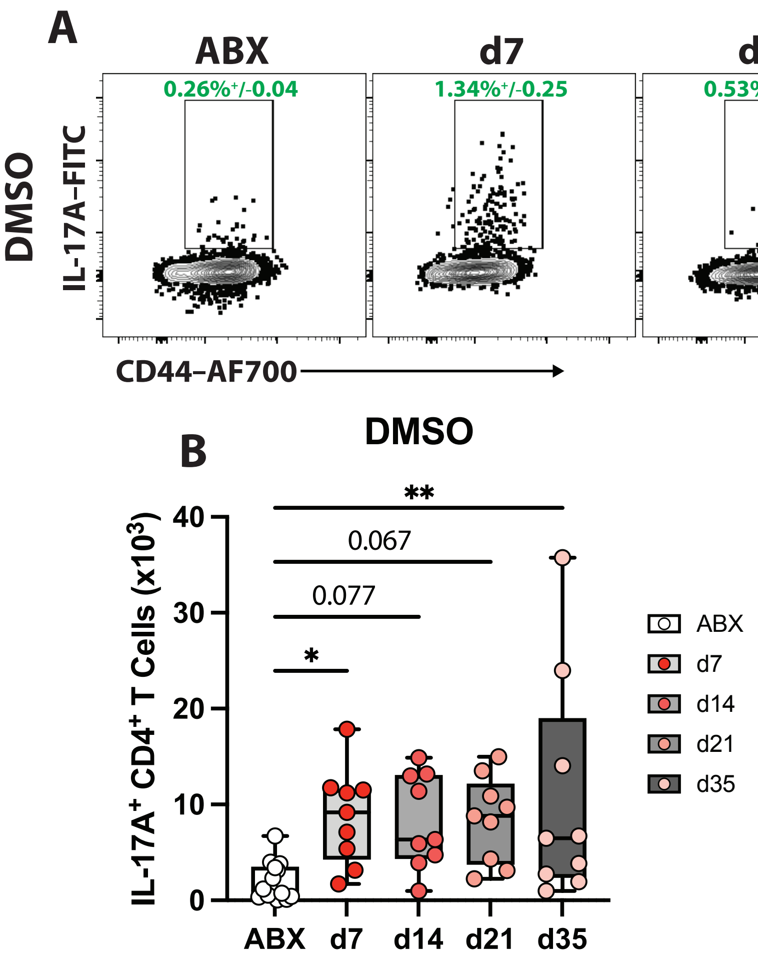

**Supplemental Figure 6**

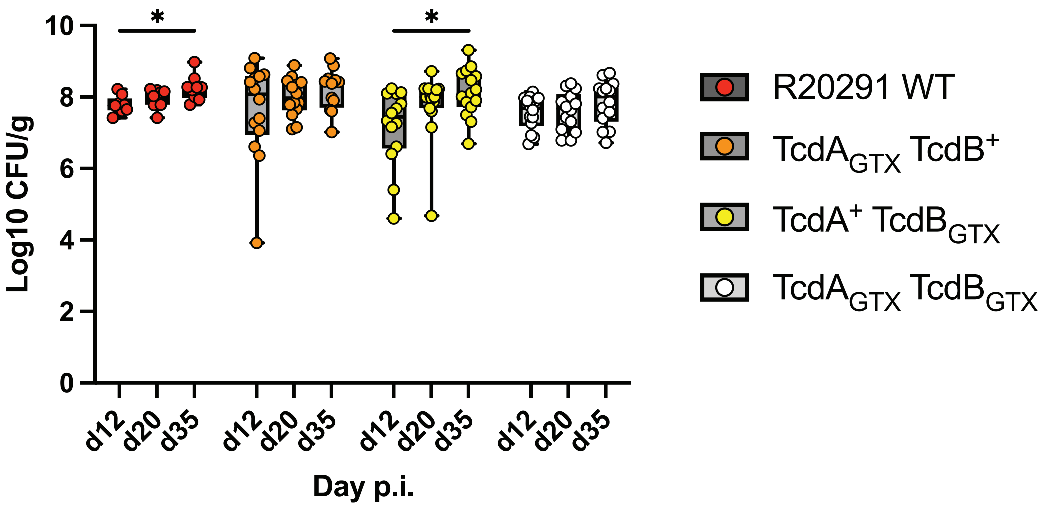

**Supplemental Figure 7**

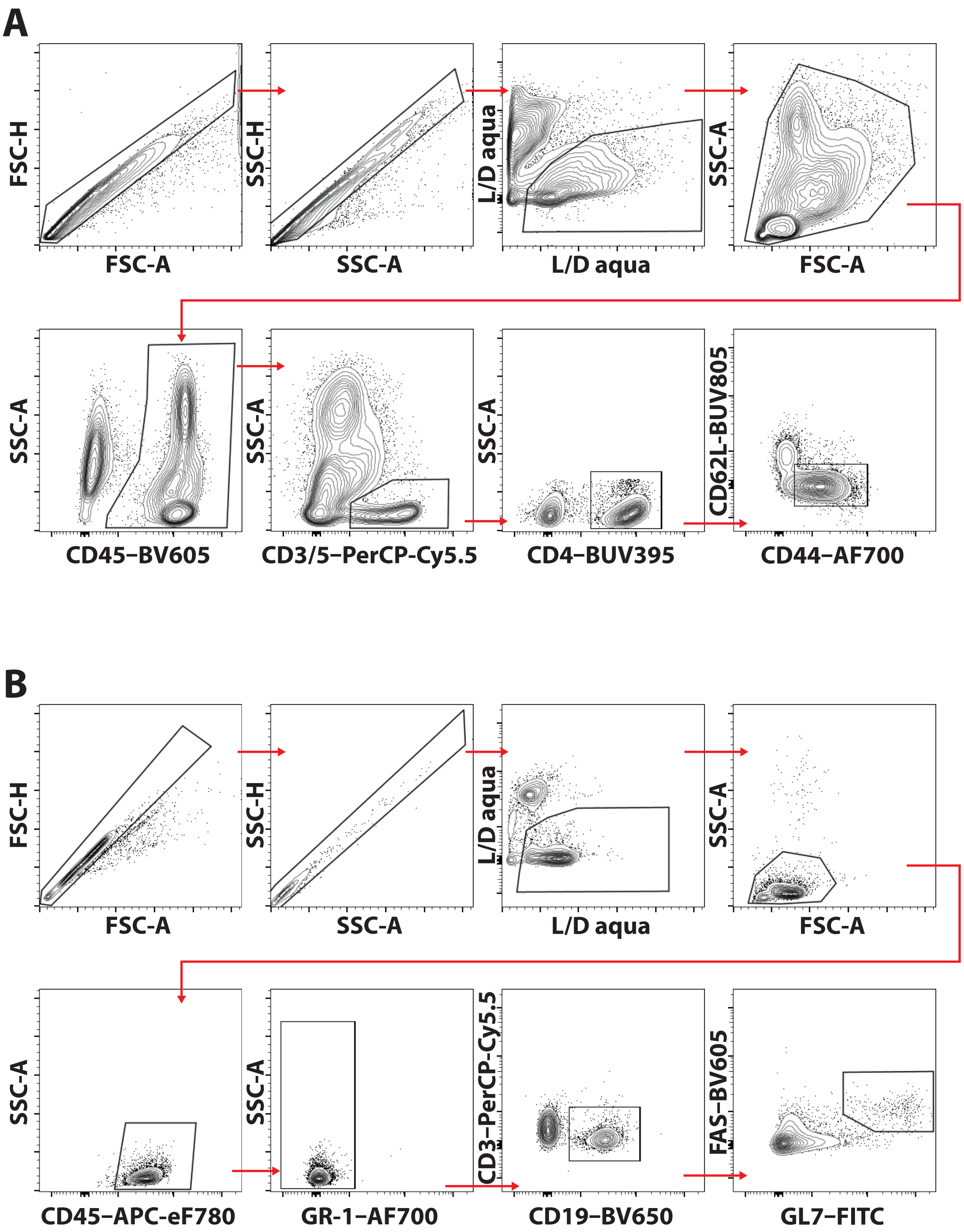

### Supplemental Table 1

**TcdA - CROP**

| Initial Final | Peptide # | Peptide Sequence |
|---|---|---|
| 1832 - 1846 | TcdA_001 | GLININNSLFYFDPI |
| 1836 - 1850 | TcdA_002 | INNSLFYFDPIESNL |
| 1840 - 1854 | TcdA_003 | LFYFDPIESNLVTGW |
| 1844 - 1858 | TcdA_004 | DPIESNLVTGWQTIN |
| 1848 - 1862 | TcdA_005 | SNLVTGWQTINGKKY |
| 1852 - 1866 | TcdA_006 | TGWQTINGKKYYFDI |
| 1856 - 1870 | TcdA_007 | TINGKKYYFDINTGA |
| 1860 - 1874 | TcdA_008 | KKYYFDINTGAASTS |
| 1864 - 1878 | TcdA_009 | FDINTGAASTSYKII |
| 1868 - 1882 | TcdA_010 | TGAASTSYKIINGKH |
| 1872 - 1886 | TcdA_011 | STSYKIINGKHFYFN |
| 1876 - 1890 | TcdA_012 | KIINGKHFYFNNNGV |
| 1880 - 1894 | TcdA_013 | GKHFYFNNNGVMQLG |
| 1884 - 1898 | TcdA_014 | YFNNNGVMQLGVFKG |
| 1888 - 1902 | TcdA_015 | NGVMQLGVFKGPDGF |
| 1892 - 1906 | TcdA_016 | QLGVFKGPDGFEYFA |
| 1896 - 1910 | TcdA_017 | FKGPDGFEYFAPANT |
| 1900 - 1914 | TcdA_018 | DGFEYFAPANTQNNN |
| 1904 - 1918 | TcdA_019 | YFAPANTQNNNIEGQ |
| 1908 - 1922 | TcdA_020 | ANTQNNNIEGQAIVY |
| 1912 - 1926 | TcdA_021 | NNNIEGQAIVYQSKF |
| 1916 - 1930 | TcdA_022 | EGQAIVYQSKFLTLN |
| 1920 - 1934 | TcdA_023 | IVYQSKFLTLNGKKY |
| 1924 - 1938 | TcdA_024 | SKFLTLNGKKYYFDN |
| 1928 - 1942 | TcdA_025 | TLNGKKYYFDNDSKA |
| 1932 - 1946 | TcdA_026 | KKYYFDNDSKAVTGW |
| 1936 - 1950 | TcdA_027 | FDNDSKAVTGWRIIN |
| 1940 - 1954 | TcdA_028 | SKAVTGWRIINNEKY |
| 1944 - 1958 | TcdA_029 | TGWRIINNEKYYFNP |
| 1948 - 1962 | TcdA_030 | IINNEKYYFNPNNAI |
| 1952 - 1966 | TcdA_031 | EKYYFNPNNAIAAVG |
| 1956 - 1970 | TcdA_032 | FNPNNAIAAVGLQVI |
| 1960 - 1974 | TcdA_033 | NAIAAVGLQVIDNNK |
| 1964 - 1978 | TcdA_034 | AVGLQVIDNNKYYFN |
| 1968 - 1982 | TcdA_035 | QVIDNNKYYFNPDTA |
| 1972 - 1986 | TcdA_036 | NNKYYFNPDTAIISK |
| 1976 - 1990 | TcdA_037 | YFNPDTAIISKGWQT |
| 1980 - 1994 | TcdA_038 | DTAIISKGWQTVNGS |
| 1984 - 1998 | TcdA_039 | ISKGWQTVNGSRYYF |
| 1988 - 2002 | TcdA_040 | WQTVNGSRYYFDTDT |
| 1992 - 2006 | TcdA_041 | NGSRYYFDTDTAIAF |
| 1996 - 2010 | TcdA_042 | YYFDTDTAIAFNGYK |
| 2000 - 2014 | TcdA_043 | TDTAIAFNGYKTIDG |
| 2004 - 2018 | TcdA_044 | IAFNGYKTIDGKHFY |
| 2008 - 2022 | TcdA_045 | GYKTIDGKHFYFDSD |
| 2012 - 2026 | TcdA_046 | IDGKHFYFDSDCVVK |
| 2016 - 2030 | TcdA_047 | HFYFDSDCVVKIGVF |
| 2020 - 2034 | TcdA_048 | DSDCVVKIGVFSGSN |
| 2024 - 2038 | TcdA_049 | VVKIGVFSGSNGFEY |
| 2028 - 2042 | TcdA_050 | GVFSGSNGFEYFAPA |
| 2032 - 2046 | TcdA_051 | GSNGFEYFAPANTYN |
| 2036 - 2050 | TcdA_052 | FEYFAPANTYNNNIE |
| 2040 - 2054 | TcdA_053 | APANTYNNNIEGQAI |
| 2044 - 2058 | TcdA_054 | TYNNNIEGQAIVYQS |
| 2048 - 2062 | TcdA_055 | NIEGQAIVYQSKFLT |
| 2052 - 2066 | TcdA_056 | QAIVYQSKFLTLNGK |
| 2056 - 2070 | TcdA_057 | YQSKFLTLNGKKYYF |
| 2060 - 2074 | TcdA_058 | FLTLNGKKYYFDNNS |
| 2064 - 2078 | TcdA_059 | NGKKYYFDNNSKAVT |
| 2068 - 2082 | TcdA_060 | YYFDNNSKAVTGWQT |
| 2072 - 2086 | TcdA_061 | NNSKAVTGWQTIDSK |
| 2076 - 2090 | TcdA_062 | AVTGWQTIDSKKYYF |
| 2080 - 2094 | TcdA_063 | WQTIDSKKYYFNTNT |
| 2084 - 2098 | TcdA_064 | DSKKYYFNTNTAEAA |
| 2088 - 2102 | TcdA_065 | YYFNTNTAEAATGWQ |
| 2092 - 2106 | TcdA_066 | TNTAEAATGWQTIDG |
| 2096 - 2110 | TcdA_067 | EAATGWQTIDGKKYY |
| 2100 - 2114 | TcdA_068 | GWQTIDGKKYYFNTN |
| 2104 - 2118 | TcdA_069 | IDGKKYYFNTNTAEA |
| 2108 - 2122 | TcdA_070 | KYYFNTNTAEAATGW |
| 2112 - 2126 | TcdA_071 | NTNTAEAATGWQTID |
| 2116 - 2130 | TcdA_072 | AEAATGWQTIDGKKY |
| 2120 - 2134 | TcdA_073 | TGWQTIDGKKYYFNT |
| 2124 - 2138 | TcdA_074 | TIDGKKYYFNTNTSI |
| 2128 - 2142 | TcdA_075 | KKYYFNTNTSIASTG |
| 2132 - 2146 | TcdA_076 | FNTNTSIASTGYTII |
| 2136 - 2150 | TcdA_077 | TSIASTGYTIINGKY |
| 2140 - 2154 | TcdA_078 | STGYTIINGKYFYFN |
| 2144 - 2158 | TcdA_079 | TIINGKYFYFNTDGI |
| 2148 - 2162 | TcdA_080 | GKYFYFNTDGIMQIG |
| 2152 - 2166 | TcdA_081 | YFNTDGIMQIGVFKV |
| 2156 - 2170 | TcdA_082 | DGIMQIGVFKVPNGF |
| 2160 - 2174 | TcdA_083 | QIGVFKVPNGFEYFA |
| 2164 - 2178 | TcdA_084 | FKVPNGFEYFAPANT |
| 2168 - 2182 | TcdA_085 | NGFEYFAPANTHNNN |
| 2172 - 2186 | TcdA_086 | YFAPANTHNNNIEGQ |
| 2176 - 2190 | TcdA_087 | ANTHNNNIEGQAILY |
| 2180 - 2194 | TcdA_088 | NNNIEGQAILYQNKF |
| 2184 - 2198 | TcdA_089 | EGQAILYQNKFLTLN |
| 2188 - 2202 | TcdA_090 | ILYQNKFLTLNGKKY |
| 2192 - 2206 | TcdA_091 | NKFLTLNGKKYYFGS |
| 2196 - 2210 | TcdA_092 | TLNGKKYYFGSDSKA |
| 2200 - 2214 | TcdA_093 | KKYYFGSDSKAITGW |
| 2204 - 2218 | TcdA_094 | FGSDSKAITGWQTID |
| 2208 - 2222 | TcdA_095 | SKAITGWQTIDGKKY |
| 2212 - 2226 | TcdA_096 | TGWQTIDGKKYYFNP |
| 2216 - 2230 | TcdA_097 | TIDGKKYYFNPNNAI |
| 2220 - 2234 | TcdA_098 | KKYYFNPNNAIAATH |
| 2224 - 2238 | TcdA_099 | FNPNNAIAATHLCTI |
| 2228 - 2242 | TcdA_100 | NAIAATHLCTINNDK |
| 2232 - 2246 | TcdA_101 | ATHLCTINNDKYYFS |
| 2236 - 2250 | TcdA_102 | CTINNDKYYFSYDGI |
| 2240 - 2254 | TcdA_103 | NDKYYFSYDGILQNG |
| 2244 - 2258 | TcdA_104 | YFSYDGILQNGYITI |
| 2248 - 2262 | TcdA_105 | DGILQNGYITIERNN |
| 2252 - 2266 | TcdA_106 | QNGYITIERNNFYFD |
| 2256 - 2270 | TcdA_107 | ITIERNNFYFDANNE |
| 2260 - 2274 | TcdA_108 | RNNFYFDANNESKMV |
| 2264 - 2278 | TcdA_109 | YFDANNESKMVTGVF |

**TcdA - CROP**

| Initial Final | Peptide # | Peptide Sequence |
|---|---|---|
| 2268 - 2282 | TcdA_110 | NNESKMVTGVFKGPN |
| 2272 - 2286 | TcdA_111 | KMVTGVFKGPNGFEY |
| 2276 - 2290 | TcdA_112 | GVFKGPNGFEYFAPA |
| 2280 - 2294 | TcdA_113 | GPNGFEYFAPANTHN |
| 2284 - 2298 | TcdA_114 | FEYFAPANTHNNNIE |
| 2288 - 2302 | TcdA_115 | APANTHNNNIEGQAI |
| 2292 - 2306 | TcdA_116 | THNNNIEGQAIVYQN |
| 2296 - 2310 | TcdA_117 | NIEGQAIVYQNKFLT |
| 2300 - 2314 | TcdA_118 | QAIVYQNKFLTLNGK |
| 2304 - 2318 | TcdA_119 | YQNKFLTLNGKKYYF |
| 2308 - 2322 | TcdA_120 | FLTLNGKKYYFDNDS |
| 2312 - 2326 | TcdA_121 | NGKKYYFDNDSKAVT |
| 2316 - 2330 | TcdA_122 | YYFDNDSKAVTGWQT |
| 2320 - 2334 | TcdA_123 | NDSKAVTGWQTIDSK |
| 2324 - 2338 | TcdA_124 | AVTGWQTIDSKKYYF |
| 2328 - 2342 | TcdA_125 | WQTIDSKKYYFNLNT |
| 2332 - 2346 | TcdA_126 | DSKKYYFNLNTAVAV |
| 2336 - 2350 | TcdA_127 | YYFNLNTAVAVTGWQ |
| 2340 - 2354 | TcdA_128 | LNTAVAVTGWQTIDG |
| 2344 - 2358 | TcdA_129 | VAVTGWQTIDGEKYY |
| 2348 - 2362 | TcdA_130 | GWQTIDGEKYYFNLN |
| 2352 - 2366 | TcdA_131 | IDGEKYYFNLNTAEA |
| 2356 - 2370 | TcdA_132 | KYYFNLNTAEAATGW |
| 2360 - 2374 | TcdA_133 | NLNTAEAATGWQTID |
| 2364 - 2378 | TcdA_134 | AEAATGWQTIDGKRY |
| 2368 - 2382 | TcdA_135 | TGWQTIDGKRYYFNT |
| 2372 - 2386 | TcdA_136 | TIDGKRYYFNTNTYI |
| 2376 - 2390 | TcdA_137 | KRYYFNTNTYIASTG |
| 2380 - 2394 | TcdA_138 | FNTNTYIASTGYTII |
| 2384 - 2398 | TcdA_139 | TYIASTGYTIINGKH |
| 2388 - 2402 | TcdA_140 | STGYTIINGKHFYFN |
| 2392 - 2406 | TcdA_141 | TIINGKHFYFNTDGI |
| 2396 - 2410 | TcdA_142 | GKHFYFNTDGIMQIG |
| 2400 - 2414 | TcdA_143 | YFNTDGIMQIGVFKG |
| 2404 - 2418 | TcdA_144 | DGIMQIGVFKGPDGF |
| 2408 - 2422 | TcdA_145 | QIGVFKGPDGFEYFA |
| 2412 - 2426 | TcdA_146 | FKGPDGFEYFAPANT |
| 2416 - 2430 | TcdA_147 | DGFEYFAPANTHNNN |
| 2420 - 2434 | TcdA_148 | YFAPANTHNNNIEGQ |
| 2424 - 2438 | TcdA_149 | ANTHNNNIEGQAILY |
| 2428 - 2442 | TcdA_150 | NNNIEGQAILYQNKF |
| 2432 - 2446 | TcdA_151 | EGQAILYQNKFLTLN |
| 2436 - 2450 | TcdA_152 | ILYQNKFLTLNGKKY |
| 2440 - 2454 | TcdA_153 | NKFLTLNGKKYYFGS |
| 2444 - 2458 | TcdA_154 | TLNGKKYYFGSDSKA |
| 2448 - 2462 | TcdA_155 | KKYYFGSDSKAVTGL |
| 2452 - 2466 | TcdA_156 | FGSDSKAVTGLRTID |
| 2456 - 2470 | TcdA_157 | SKAVTGLRTIDGKKY |
| 2460 - 2474 | TcdA_158 | TGLRTIDGKKYYFNT |
| 2464 - 2478 | TcdA_159 | TIDGKKYYFNTNTAV |
| 2468 - 2482 | TcdA_160 | KKYYFNTNTAVAVTG |
| 2472 - 2486 | TcdA_161 | FNTNTAVAVTGWQTI |
| 2476 - 2490 | TcdA_162 | TAVAVTGWQTINGKK |
| 2480 - 2494 | TcdA_163 | VTGWQTINGKKYYFN |
| 2484 - 2498 | TcdA_164 | QTINGKKYYFNTNTY |
| 2488 - 2502 | TcdA_165 | GKKYYFNTNTYIAST |
| 2492 - 2506 | TcdA_166 | YFNTNTYIASTGYTI |
| 2496 - 2510 | TcdA_167 | NTYIASTGYTIISGK |
| 2500 - 2514 | TcdA_168 | ASTGYTIISGKHFYF |
| 2504 - 2518 | TcdA_169 | YTIISGKHFYFNTDG |
| 2508 - 2522 | TcdA_170 | SGKHFYFNTDGIMQI |
| 2512 - 2526 | TcdA_171 | FYFNTDGIMQIGVFK |
| 2516 - 2530 | TcdA_172 | TDGIMQIGVFKGPDG |
| 2520 - 2534 | TcdA_173 | MQIGVFKGPDGFEYF |
| 2524 - 2538 | TcdA_174 | VFKGPDGFEYFAPAN |
| 2528 - 2542 | TcdA_175 | PDGFEYFAPANTDAN |
| 2532 - 2546 | TcdA_176 | EYFAPANTDANNIEG |
| 2536 - 2550 | TcdA_177 | PANTDANNIEGQAIR |
| 2540 - 2554 | TcdA_178 | DANNIEGQAIRYQNR |
| 2544 - 2558 | TcdA_179 | IEGQAIRYQNRFLYL |
| 2548 - 2562 | TcdA_180 | AIRYQNRFLYLHDNI |
| 2552 - 2566 | TcdA_181 | QNRFLYLHDNIYYFG |
| 2556 - 2570 | TcdA_182 | LYLHDNIYYFGNDSK |
| 2560 - 2574 | TcdA_183 | DNIYYFGNDSKAATG |
| 2564 - 2578 | TcdA_184 | YFGNDSKAATGWATI |
| 2568 - 2582 | TcdA_185 | DSKAATGWATIDGNR |
| 2572 - 2586 | TcdA_186 | ATGWATIDGNRYYFE |
| 2576 - 2590 | TcdA_187 | ATIDGNRYYFEPNTA |
| 2580 - 2594 | TcdA_188 | GNRYYFEPNTAMGAN |
| 2584 - 2598 | TcdA_189 | YFEPNTAMGANGYKT |
| 2588 - 2602 | TcdA_190 | NTAMGANGYKTIDNK |
| 2592 - 2606 | TcdA_191 | GANGYKTIDNKNFYF |
| 2596 - 2610 | TcdA_192 | YKTIDNKNFYFRNGL |
| 2600 - 2614 | TcdA_193 | DNKNFYFRNGLPQIG |
| 2604 - 2618 | TcdA_194 | FYFRNGLPQIGVFKG |
| 2608 - 2622 | TcdA_195 | NGLPQIGVFKGPNGF |
| 2612 - 2626 | TcdA_196 | QIGVFKGPNGFEYFA |
| 2616 - 2630 | TcdA_197 | FKGPNGFEYFAPANT |
| 2620 - 2634 | TcdA_198 | NGFEYFAPANTDANN |
| 2624 - 2638 | TcdA_199 | YFAPANTDANNIDGQ |
| 2628 - 2642 | TcdA_200 | ANTDANNIDGQAIRY |
| 2632 - 2646 | TcdA_201 | ANNIDGQAIRYQNRF |
| 2636 - 2650 | TcdA_202 | DGQAIRYQNRFLHLL |
| 2640 - 2654 | TcdA_203 | IRYQNRFLHLLGKIY |
| 2644 - 2658 | TcdA_204 | NRFLHLLGKIYYFGN |
| 2648 - 2662 | TcdA_205 | HLLGKIYYFGNNSKA |
| 2652 - 2666 | TcdA_206 | KIYYFGNNSKAVTGW |
| 2656 - 2670 | TcdA_207 | FGNNSKAVTGWQTIN |
| 2660 - 2674 | TcdA_208 | SKAVTGWQTINSKVY |
| 2664 - 2678 | TcdA_209 | TGWQTINSKVYYFMP |
| 2668 - 2682 | TcdA_210 | TINSKVYYFMPDTAM |
| 2672 - 2686 | TcdA_211 | KVYYFMPDTAMAAAG |
| 2676 - 2690 | TcdA_212 | FMPDTAMAAAGGLFE |
| 2680 - 2694 | TcdA_213 | TAMAAAGGLFEIDGV |
| 2684 - 2698 | TcdA_214 | AAGGLFEIDGVIYFF |
| 2688 - 2702 | TcdA_215 | LFEIDGVIYFFGVDG |
| 2692 - 2706 | TcdA_216 | DGVIYFFGVDGVKAP |
| 2696 - 2710 | TcdA_217 | YFFGVDGVKAPGIYG |

**TcdB - CROP**

| Initial Final | Peptide # | Peptide Sequence |
|---|---|---|
| 1836 - 1850 | TcdB_001 | VYINDSLYYFKPPIK |
| 1837 - 1851 | TcdB_002 | DSLYYFKPPIKNLIT |
| 1838 - 1852 | TcdB_003 | YFKPPIKNLITGFTT |
| 1839 - 1853 | TcdB_004 | PIKNLITGFTTIGDD |
| 1840 - 1854 | TcdB_005 | LITGFTTIGDDKYYF |
| 1841 - 1855 | TcdB_006 | FTTIGDDKYYFNPDN |
| 1842 - 1856 | TcdB_007 | GDDKYYFNPDNGGAA |
| 1843 - 1857 | TcdB_008 | YYFNPDNGGAASVGE |
| 1844 - 1858 | TcdB_009 | PDNGGAASVGETIID |
| 1845 - 1859 | TcdB_010 | GAASVGETIIDGKNY |
| 1846 - 1860 | TcdB_011 | VGETIIDGKNYYFSQ |
| 1847 - 1861 | TcdB_012 | IIDGKNYYFSQNGVL |
| 1848 - 1862 | TcdB_013 | KNYYFSQNGVLQTGV |
| 1849 - 1863 | TcdB_014 | FSQNGVLQTGVFSTE |
| 1850 - 1864 | TcdB_015 | GVLQTGVFSTEDGFK |
| 1851 - 1865 | TcdB_016 | TGVFSTEDGFKYFAP |
| 1852 - 1866 | TcdB_017 | STEDGFKYFAPADTL |
| 1853 - 1867 | TcdB_018 | GFKYFAPADTLDENL |
| 1854 - 1868 | TcdB_019 | FAPADTLDENLEGEA |
| 1855 - 1869 | TcdB_020 | DTLDENLEGEAIDFT |
| 1856 - 1870 | TcdB_021 | ENLEGEAIDFTGKLT |
| 1857 - 1871 | TcdB_022 | GEAIDFTGKLTIDEN |
| 1858 - 1872 | TcdB_023 | DFTGKLTIDENVYYF |
| 1859 - 1873 | TcdB_024 | KLTIDENVYYFGDNY |
| 1860 - 1874 | TcdB_025 | DENVYYFGDNYRAAI |
| 1861 - 1875 | TcdB_026 | YYFGDNYRAAIEWQT |
| 1862 - 1876 | TcdB_027 | DNYRAAIEWQTLDDE |
| 1863 - 1877 | TcdB_028 | AAIEWQTLDDEVYYF |
| 1864 - 1878 | TcdB_029 | WQTLDDEVYYFSTDT |
| 1865 - 1879 | TcdB_030 | DDEVYYFSTDTGRAF |
| 1866 - 1880 | TcdB_031 | YYFSTDTGRAFKGLN |
| 1867 - 1881 | TcdB_032 | TDTGRAFKGLNQIGD |
| 1868 - 1882 | TcdB_033 | RAFKGLNQIGDDKFY |
| 1869 - 1883 | TcdB_034 | GLNQIGDDKFYFNSD |
| 1870 - 1884 | TcdB_035 | IGDDKFYFNSDGIMQ |
| 1871 - 1885 | TcdB_036 | KFYFNSDGIMQKGFV |
| 1872 - 1886 | TcdB_037 | NSDGIMQKGFVNIND |
| 1873 - 1887 | TcdB_038 | IMQKGFVNINDKTFY |
| 1874 - 1888 | TcdB_039 | GFVNINDKTFYFDDS |
| 1875 - 1889 | TcdB_040 | INDKTFYFDDSGVMK |
| 1876 - 1890 | TcdB_041 | TFYFDDSGVMKSGYT |
| 1877 - 1891 | TcdB_042 | DDSGVMKSGYTEIDG |
| 1878 - 1892 | TcdB_043 | VMKSGYTEIDGKYFY |
| 1879 - 1893 | TcdB_044 | GYTEIDGKYFYFAEN |
| 1880 - 1894 | TcdB_045 | IDGKYFYFAENGEMQ |
| 1881 - 1895 | TcdB_046 | YFYFAENGEMQIGVF |
| 1882 - 1896 | TcdB_047 | AENGEMQIGVFNTAD |
| 1883 - 1897 | TcdB_048 | EMQIGVFNTADGFKY |
| 1884 - 1898 | TcdB_049 | GVFNTADGFKYFAHH |
| 1885 - 1899 | TcdB_050 | TADGFKYFAHHDEDL |
| 1886 - 1900 | TcdB_051 | FKYFAHHDEDLGNEE |
| 1887 - 1901 | TcdB_052 | AHHDEDLGNEEGEAL |
| 1888 - 1902 | TcdB_053 | EDLGNEEGEALSYSG |
| 1889 - 1903 | TcdB_054 | NEEGEALSYSGILNF |
| 1890 - 1904 | TcdB_055 | EALSYSGILNFNNKI |
| 1891 - 1905 | TcdB_056 | YSGILNFNNKIYYFD |
| 1892 - 1906 | TcdB_057 | LNFNNKIYYFDDSFT |
| 1893 - 1907 | TcdB_058 | NKIYYFDDSFTAVVG |
| 1894 - 1908 | TcdB_059 | YFDDSFTAVVGWKDL |
| 1895 - 1909 | TcdB_060 | SFTAVVGWKDLEDGS |
| 1896 - 1910 | TcdB_061 | VVGWKDLEDGSKYYF |
| 1897 - 1911 | TcdB_062 | KDLEDGSKYYFDEDT |
| 1898 - 1912 | TcdB_063 | DGSKYYFDEDTAEAY |
| 1899 - 1913 | TcdB_064 | YYFDEDTAEAYIGIS |
| 1900 - 1914 | TcdB_065 | EDTAEAYIGISIIND |

**TcdB - CROP**

| Initial Final | Peptide # | Peptide Sequence |
|---|---|---|
| 1901 - 1915 | TcdB_066 | EAYIGISIINDGKYY |
| 1902 - 1916 | TcdB_067 | GISIINDGKYYFNDS |
| 1903 - 1917 | TcdB_068 | INDGKYYFNDSGIMQ |
| 1904 - 1918 | TcdB_069 | KYYFNDSGIMQIGFV |
| 1905 - 1919 | TcdB_070 | NDSGIMQIGFVTINN |
| 1906 - 1920 | TcdB_071 | IMQIGFVTINNEVFY |
| 1907 - 1921 | TcdB_072 | GFVTINNEVFYFSDS |
| 1908 - 1922 | TcdB_073 | INNEVFYFSDSGIVE |
| 1909 - 1923 | TcdB_074 | VFYFSDSGIVESGMQ |
| 1910 - 1924 | TcdB_075 | SDSGIVESGMQNIDD |
| 1911 - 1925 | TcdB_076 | IVESGMQNIDDNYFY |
| 1912 - 1926 | TcdB_077 | GMQNIDDNYFYIDEN |
| 1913 - 1927 | TcdB_078 | IDDNYFYIDENGLVQ |
| 1914 - 1928 | TcdB_079 | YFYIDENGLVQIGVF |
| 1915 - 1929 | TcdB_080 | DENGLVQIGVFDTSD |
| 1916 - 1930 | TcdB_081 | LVQIGVFDTSDGYKY |
| 1917 - 1931 | TcdB_082 | GVFDTSDGYKYFAPA |
| 1918 - 1932 | TcdB_083 | TSDGYKYFAPANTVN |
| 1919 - 1933 | TcdB_084 | YKYFAPANTVNDNIY |
| 1920 - 1934 | TcdB_085 | APANTVNDNIYGQAV |
| 1921 - 1935 | TcdB_086 | TVNDNIYGQAVEYSG |
| 1922 - 1936 | TcdB_087 | NIYGQAVEYSGLVRV |
| 1923 - 1937 | TcdB_088 | QAVEYSGLVRVGEDV |
| 1924 - 1938 | TcdB_089 | YSGLVRVGEDVYYFG |
| 1925 - 1939 | TcdB_090 | VRVGEDVYYFGETYT |
| 1926 - 1940 | TcdB_091 | EDVYYFGETYTIETG |
| 1927 - 1941 | TcdB_092 | YFGETYTIETGWIYD |
| 1928 - 1942 | TcdB_093 | TYTIETGWIYDMENE |
| 1929 - 1943 | TcdB_094 | ETGWIYDMENESDKY |
| 1930 - 1944 | TcdB_095 | IYDMENESDKYYFDP |
| 1931 - 1945 | TcdB_096 | ENESDKYYFDPETKK |
| 1932 - 1946 | TcdB_097 | DKYYFDPETKKAYKG |
| 1933 - 1947 | TcdB_098 | FDPETKKAYKGINVI |
| 1934 - 1948 | TcdB_099 | TKKAYKGINVIDDIK |
| 1935 - 1949 | TcdB_100 | YKGINVIDDIKYYFD |
| 1936 - 1950 | TcdB_101 | NVIDDIKYYFDENGI |
| 1937 - 1951 | TcdB_102 | DIKYYFDENGIMRTG |
| 1938 - 1952 | TcdB_103 | YFDENGIMRTGLITF |
| 1939 - 1953 | TcdB_104 | NGIMRTGLITFEDNH |
| 1940 - 1954 | TcdB_105 | RTGLITFEDNHYYFN |
| 1941 - 1955 | TcdB_106 | ITFEDNHYYFNEDGI |
| 1942 - 1956 | TcdB_107 | DNHYYFNEDGIMQYG |
| 1943 - 1957 | TcdB_108 | YFNEDGIMQYGYLNI |
| 1944 - 1958 | TcdB_109 | DGIMQYGYLNIEDKT |
| 1945 - 1959 | TcdB_110 | QYGYLNIEDKTFYFS |
| 1946 - 1960 | TcdB_111 | LNIEDKTFYFSEDGI |
| 1947 - 1961 | TcdB_112 | DKTFYFSEDGIMQIG |
| 1948 - 1962 | TcdB_113 | YFSEDGIMQIGVFNT |
| 1949 - 1963 | TcdB_114 | DGIMQIGVFNTPDGF |
| 1950 - 1964 | TcdB_115 | QIGVFNTPDGFKYFA |
| 1951 - 1965 | TcdB_116 | FNTPDGFKYFAHQNT |
| 1952 - 1966 | TcdB_117 | DGFKYFAHQNTLDEN |
| 1953 - 1967 | TcdB_118 | YFAHQNTLDENFEGE |
| 1954 - 1968 | TcdB_119 | QNTLDENFEGESINY |
| 1955 - 1969 | TcdB_120 | DENFEGESINYTGWL |
| 1956 - 1970 | TcdB_121 | EGESINYTGWLDLDE |
| 1957 - 1971 | TcdB_122 | INYTGWLDLDEKRYY |
| 1958 - 1972 | TcdB_123 | GWLDLDEKRYYFTDE |
| 1959 - 1973 | TcdB_124 | LDEKRYYFTDEYIAA |
| 1960 - 1974 | TcdB_125 | RYYFTDEYIAATGSV |
| 1961 - 1975 | TcdB_126 | TDEYIAATGSVIIDG |
| 1962 - 1976 | TcdB_127 | IAATGSVIIDGEEYY |
| 1963 - 1977 | TcdB_128 | GSVIIDGEEYYFDPD |
| 1964 - 1978 | TcdB_129 | IDGEEYYFDPDTAQL |
| 1965 - 1979 | TcdB_130 | EYYFDPDTAQLVISE |
